# Acute immune modulation with poly-salicylic acid particles ameliorates pain and structural damage in post-traumatic osteoarthritis

**DOI:** 10.1101/2025.11.21.689790

**Authors:** Michael LaRue Felder, Lindsey Lammlin, Adrienne A. Giannone, M. Valentina Guevara, Scarlet C. Howser, Isabelle J. Smith, Omolola Eniola-Adefeso, Tristan Maerz

**Affiliations:** University of Michigan, Ann Arbor, MI, United States; ETH Zürich, Zürich, Switzerland; University of Illinois Chicago, Chicago, IL, United States

## Abstract

Joint inflammation is a hallmark of post-traumatic osteoarthritis (PTOA) progression and a recognized driver of articular destruction and symptoms. Despite its known pathological role, inflammation has not been successfully targeted to treat PTOA. With the hypothesis that blocking the acute influx of systemically-derived immune cells can mitigate injury-induced inflammation and downstream PTOA disease severity, we targeted immune cell recruitment via systemically-administered poly salicylic acid (PolySA) particles. This formulation targets immune cells in circulation, namely neutrophils and monocytes, to inhibit their vascular extravasation into injured tissue. Employing a murine joint injury model, we show that PolySA particles reduced neutrophil and monocyte recruitment to the synovium by >50% when administered acutely after injury. Sex-specific therapeutic effects of PolySA emerged 7d post-ACLR, whereby female knee joints exhibited increased cathepsin activity and alleviation of knee hyperalgesia. Despite also observing reduced immune cell recruitment in male mice treated with PolySA, therapeutic effects were entirely absent in males. RNAseq of female synovium revealed a transcriptomic signature indicative of accelerated immune resolution and matrix remodeling in PolySA-treated female mice. Analyses at a timepoint of established disease showed that PolySA-treated female mice exhibited sustained pain alleviation, reduced osteophyte formation, and decreased histopathological PTOA and synovitis severity scores. Together, these findings indicate that blocking acutely-recruited immune cells to the local joint microenvironment via systemic PolySA particle treatment is a promising therapeutic for PTOA prevention by reprogramming early injury-induced inflammation.

## Introduction

Successful disease-modifying treatment of osteoarthritis (OA) has not been achieved. Current clinical management of OA is primarily focused on analgesia and is entirely palliative, and chronic use of these analgesic strategies is associated with significant side effects.^1–3^ Total joint arthroplasty is employed at end-stage disease, at which point OA patients have had to endure decades of pain, poor mobility, and numerous medical and psychosocial comorbidities. Thus, there is a critical need for novel disease-modifying treatments that target the root cause of OA. Novel formulations that involve cell specific-targeting hold significant potential for improved therapeutic efficacy and minimization of adverse side effects.

In the context of post-traumatic OA (PTOA), which develops following joint injury and represents ∼12% of all OA cases^4^, the severe joint inflammation that develops rapidly after injury is a major trigger of disease and represents a unique opportunity for early intervention to block disease onset. Despite the ineffectiveness of RA drugs in OA^5^, extensive evidence exists for the therapeutic potential of targeting inflammation in OA and PTOA, and synovial inflammation is a well-recognized driver of OA progression and pain.^6^ Clinical studies have shown that synovial inflammation is a predictor of overall OA severity and is correlated with worsened disease outcomes.^7,8^ As macrophages are the most abundant immune cell in the OA joint, extensive prior work has focused on targeting macrophages to alleviate OA inflammation. However, attempts to ablate the injury-induced inflammatory response by complete macrophage depletion has resulted in mixed outcomes. Using the destabilization of the medial meniscus (DMM) PTOA model in macrophage Fas-induced apoptosis (MaFIA) mice, Wu, et. al., showed that macrophage depletion at the time of injury resulted in more severe synovitis, increased intra-articular neutrophil and CD3+ T cell proportions, and increased levels of pro-inflammatory cytokines in synovial fluid and serum at 9 wks post-DMM.^9^ In contrast, other work shows that macrophage depletion in MaFIA mice at established PTOA timepoints (8 and 16 wks post-DMM, 12 wks post-partial meniscectomy (PMX)) resulted in pain mitigation, which was postulated to be mediated by decreased neutrophils and pro-inflammatory macrophage content in the dorsal root ganglion (DRG), but no effect on structural damage was observed in the joint.^10^ These studies highlight the need for therapeutics that optimally modulate the nature, magnitude, and dynamics of the joint’s inflammatory response for effective disease mitigation to be achieved.

Monocytes and macrophages have been the focus of most attention when studying and targeting PTOA-related inflammation^11–13^ However, neutrophils are the most abundant immune cell in blood and are often considered the first responders to sites of trauma and inflammation, preceding monocyte infiltration.^14,15^ As such, early neutrophil activation directs the course of the subsequent immune response. Neutrophils have been identified as a disease-associated cell type in both early and late phases of OA, as well as a major contributor to the secretion of inflammatory cytokines, chemokines, and proteases that propagate inflammation and tissue destruction, making them a viable target for treatment of PTOA inflammation.^13,16^ In human OA subjects, neutrophils were shown to abundant in synovial fluid, where they produced matrix degrading neutrophil elastase and were associated with worse OA progression.^16^ Therefore, targeting the early immune cell responders to joint injury, namely neutrophils, represents a promising strategy to alter the development and progression of PTOA.

Particle-based therapeutics are a viable and emerging strategy to target immune cells. Polymeric particle formulations have been shown to interact with and reroute immune cells in circulation, providing the unique opportunity to target immune cells before their recruitment to injured tissues and to avoid off-target effects.^17^ These formulations have been employed for treating acute inflammation and autoimmune disorders, where circulating innate immune cells internalize particles to reroute them to the spleen and liver.^18–20^ One formulation of particular interest is poly salicylic acid (PolySA) microparticles for the modulation of acute neutrophilic inflammation.^19,21–23^ We have shown that PolySA microparticles degrade into salicylic acid, the active component in aspirin, a well-known, safe NSAID.^22^ Further, PolySA particles have exhibited distinct anti-inflammatory activity not observed by free salicylic acid and other non-therapeutic polymer particles.^19,23^ Thus, PolySA could be utilized for its cell-targeted and controlled release of a common NSAID for use in PTOA.

Here, we demonstrate the beneficial therapeutic effects of targeting systemically-derived immune cell influx into the joint via PolySA particle treatment. We hypothesized that rerouting of immune cells to block their intra-articular recruitment can alter PTOA development after joint injury. Our results demonstrate across multiple readouts that acute systemic administration of PolySA induces reprogramming of injury-induced joint inflammation that results in mitigated pain and blunted structural disease progression.

## Results

### Systemic PolySA particle treatment mitigates intra-articular immune cell recruitment following joint injury

To inform therapeutic timing in the context of murine PTOA, we first sought to establish the kinetics of immune cell recruitment to the synovium following joint injury. To faithfully model the immune response following joint trauma, we employed a clinically relevant, noninvasive joint injury model that permits the assessments of early timepoints due to the absence of biological confounders from surgical injury.^24^ Flow cytometric analysis of injured synovia was performed at seven timepoints acutely post-ACLR (0hr to 48hrs). We observed the time-dependent influx of neutrophils as early as 4-8 hrs post-injury, with peak neutrophil counts observed at 24hrs post-injury (**Fig. 1A**). Monocytes followed neutrophils in their influx at 8hrs following ACLR, with a relative peak at ∼28hrs post-injury (**Fig. 1B**). Peak monocyte content was approximately double of neutrophil content. Finally, synovial macrophage numbers increased from 8hrs onwards, with a distinct population shift observed by flow cytometry (**Fig. 1C**). Given the presence of resident synovial macrophages that are also known to proliferate after injury and in response to intra-articular inflammatory signals^25^, this increase in macrophage content was likely driven by both local proliferation and exogenous influx of monocytes that differentiate into mature macrophages. Statistical analysis comparing males (triangle) to females (circle) indicated no difference in immune cell infiltration at any time point (**Fig. 1A-C**).

**Figure 1:**
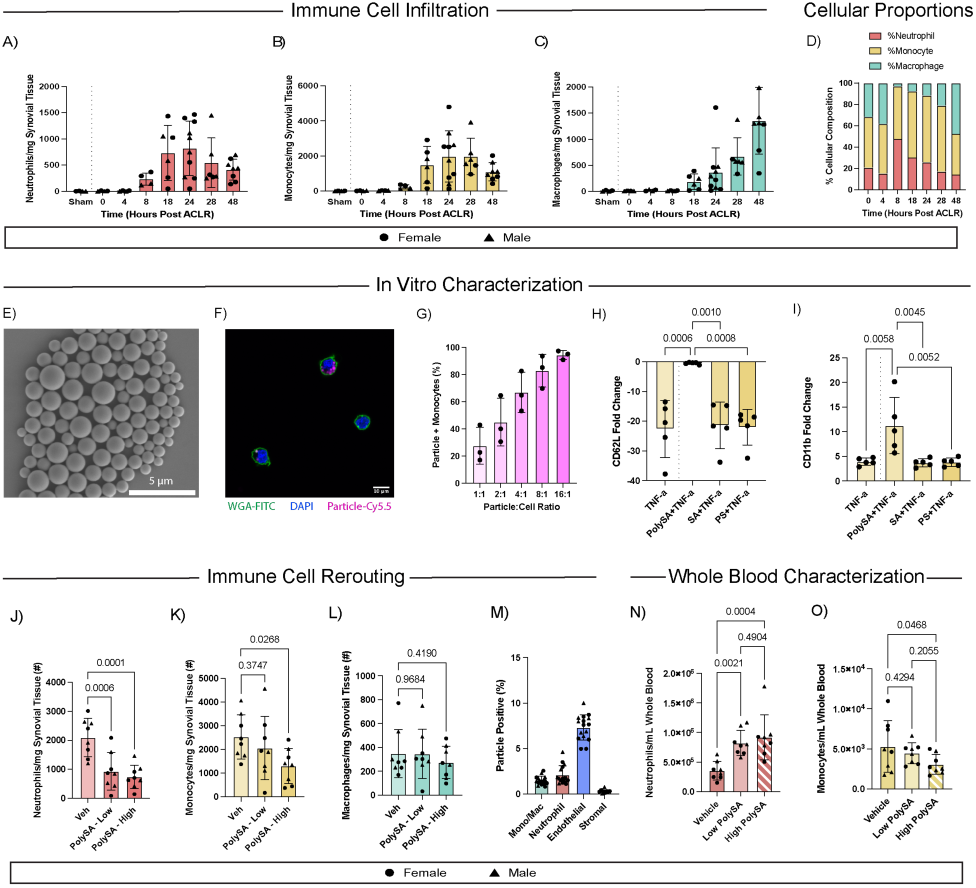
Immune cell recruitment to the synovium. Influx of (A) neutrophils, (B), monocytes, and (C) macrophages into the synovium from 0hrs to 48hrs post-ACLR as measured by flow cytometry (n≥6; male (triangle) and female (circle) depicted on graph). Welch’s t-test was used to determine differences between sexes at each timepoint. D) Immune cell proportions in the synovium during acute immune activation (0hrs to 48hrs) post-ACLR. (E) Representative scanning electron microscopy image of PolySA particles. (F) Confocal microscopy image depicting human monocytes with nuclear stain (blue), cell membrane stain (green), and internalized PolySA-Cy5.5 particles (magenta). Scale bar is 10 µm. (G) Particle-positive human monocytes after incubation with PolySA-Cy5.5 particles at varying ratios (n=3 per condition). (H-I) Surface protein expression by flow cytometry of TNF-a activated human monocytes following treatment with salicylic acid, polystyrene particles, and PolySA particles, as measured by (H) CD62L and (I) CD11b. One-Way ANOVA with Tukey’s post hoc test (n=5). (J) Neutrophil, (K) monocyte, and (L) macrophage counts in the synovium at 24hrs post-ACLR following treatment with Vehicle (Veh, PBS), low-dose PolySA (2x10^8^ particles) or high-dose PolySA (4x10^8^ particles) at 12hrs post-ACLR. One-Way ANOVA with post hoc Fisher’s LSD (n=8; 4 male/4 female). Welch’s t-test used to determine differences between sexes for each treatment. (M) Percentage of Cy5.5-positive cells across synovial cell types following treatment with high-dose Cy5.5-labeled PolySA at 12 hrs post-ACLR, with analysis at 24 hrs (n=8; 4 male/4 female). Whole blood cell counts of (N) neutrophils and (O) monocytes in Vehicle, low-dose PolySA, and high-dose PolySA at 24 hours post-ACLR. One-Way ANOVA with post hoc Fisher’s LSD (n=8; 4 male/4 female).

Having characterized the time-dependent cell trafficking into the injured synovium, we next sought to modulate these cell dynamics by rerouting cells away from the site of inflammation using a systemically administered anti-inflammatory particle therapeutic – Poly Salicylic Acid (PolySA). Our group previously characterized the neutrophilic response to PolySA, demonstrating the rerouting of circulating neutrophils to the filtering organs to block their recruitment to sites of inflammation.^19,22^ Due to the large recruitment of monocytes to the injured joint and given the phagocytic capacity of monocytes, we first extended our prior studies to characterize the uptake of PolySA particles by monocytes and the impact of PolySA particles on monocyte behavior. We generated PolySA particles (**Fig. 1E**) with a monodispersed particle population (1.14 +/- 0.33 µm) (**Suppl. Fig. 1**) that had an average zeta potential of -16.5 mV, consistent with previous work.^19^ Human monocytes readily internalized PolySA particles in a dose-dependent manner, as measured by confocal microscopy and flow cytometry, respectively (**Fig. 1F-G**). PolySA-positive monocytes exhibited complete inhibition of TNF-α-induced reduction of CD62L/L-selectin, a cell-surface protein critical for leukocyte adhesion and motility, and this was not observed followed treatment with dose-equivalent free salicylic acid or inert polystyrene particles of equivalent size (**Fig 1H**). We further observed that PolySA increased the TNF-α-induced enrichment of CD11b, which was not induced by free SA or polystyrene particles (**Fig. 1I**). While CD62L and CD11b expression for PolySA-positive monocytes is somewhat unexpected, due to the fact that CD11b upregulation is indicative of monocyte activation and CD62L retention is indicative of anti-inflammatory activity, these findings nonetheless demonstrate that PolySA also targets monocytes to modulate their phenotype.

We next aimed to assess whether acute intervention with PolySA could block immune cell recruitment into the synovium of injured joints. We systemically administered a vehicle control or PolySA at 12hrs post-ACLR, the timepoint of half-maximum neutrophil influx based on our findings (**Fig. 1A**). Treatments were administered retro-orbitally, and we tested two doses of PolySA particles: 2x10^8^ particles in 100 μL (low dose) and 4x10^8^ particles in 100 μL (high dose). Flow cytometric analysis of injured synovium at 24hrs post-ACLR (i.e. 12 hrs post-treatment) demonstrated that PolySA treatment reduced the number of tissue neutrophils by 56% (*P*=0.0006) and 65% (*P*=0.0001) at the low and high dose, respectively, relative to the vehicle control (**Fig. 1J**). High dose PolySA also reduced the total number of monocytes by 48% (*P*=0.0268) relative to control (**Fig. 1K**). No effect on the total number of synovial macrophages was observed at either dose of PolySA (**Fig. 1L**). Welch’s t-tests were conducted to determine any effects of sex on immune cell numbers, and we observed no differences between male and female synovial immune cell counts in any cell type or treatment. Consistent with PolySA-mediated blockade of recruitment to the joint, we also observed higher neutrophil counts in whole blood of both low and high dose PolySA-treated mice relative to the vehicle treated control (**Fig. 1N**). Conversely, we observed lower monocyte counts in whole blood of high-dose PolySA-treated mice compared to vehicle (**Fig. 1O**).

To further confirm that PolySA-targeted immune cells exhibit mitigated recruitment to the injured joint, we treated a separate cohort of mice at 12hrs post-ACLR with Cy5.5-labeled high-dose PolySA particles (Cy5.5-PolySA) and analyzed synovial cells at 24hrs post-injury using flow cytometry. We identified a very low proportion of particle-positive cells within synovial tissue at 24 hours following administration of the high dose of PolySA (**Fig. 1M**). Less than 2% of all monocytes, macrophages, and neutrophils were Cy5.5-positive, suggesting that nearly all particle-positive immune cells were rerouted away from the injured joint, likely to the spleen or liver, consistent with our previous work in a lung injury model.^22^ We observed that ∼7% of endothelial cells and a negligible proportion (<0.5%) of stromal cells were Cy5.5-positive.

Taken together, these findings demonstrate that acute, systemic PolySA treatment effectively mitigated monocyte and neutrophil recruitment to synovium during the acute phase of inflammation following joint injury.

### Acute PolySA treatment mitigates knee hyperalgesia and alters joint inflammation in a sex-specific manner

We next sought to understand the downstream therapeutic effects of PolySA-mediated blockade of immune cell recruitment on early-stage synovitis and PTOA development. The acute phase following major joint trauma represents a distinct opportunity for clinical intervention given that most patients will seek medical attention following injury. Accordingly, we utilized a clinically-relevant therapeutic regimen by administering two acute treatments at 12hr and 48hr post-ACLR of either vehicle or PolySA (4x10^8^ particles in 100 μL), followed by evaluation of synovial immune cell recruitment, pain behavior via knee hyperalgesia assessment, molecular imaging of peri-articular cathepsin activity as a broad readout of inflammation, and histopathologic changes at 7d post-ACLR, a timepoint of robust synovial inflammation preceding PTOA development^26^ (**Fig. 2A**). Flow cytometric analysis demonstrated that PolySA-treated mice exhibited similar synovial neutrophil and macrophage content but higher monocyte content compared to vehicle-treated mice (**Suppl. Fig. 2**). Notably, monocyte content at 7d post-ACLR was very low compared to the 24-hr timepoint, and the absolute value of the observed difference was small (156 monocytes/mg synovial tissue vs 237 monocytes/mg synovial tissue on average). Whole blood cell analysis of females showed no difference in total neutrophil and monocyte counts per mL of whole blood at 7d post-ACLR (**Fig. 2C and 2E**). However, male mice showed significantly higher neutrophil counts in whole blood of PolySA-treated mice and significantly lower monocyte counts in PolySA-treated mice compared to control, indicating a longer term systemic effect in males for the 12-hr and 48-hr treatment strategy (**Fig. 2B and 2D**).

**Figure 2:**
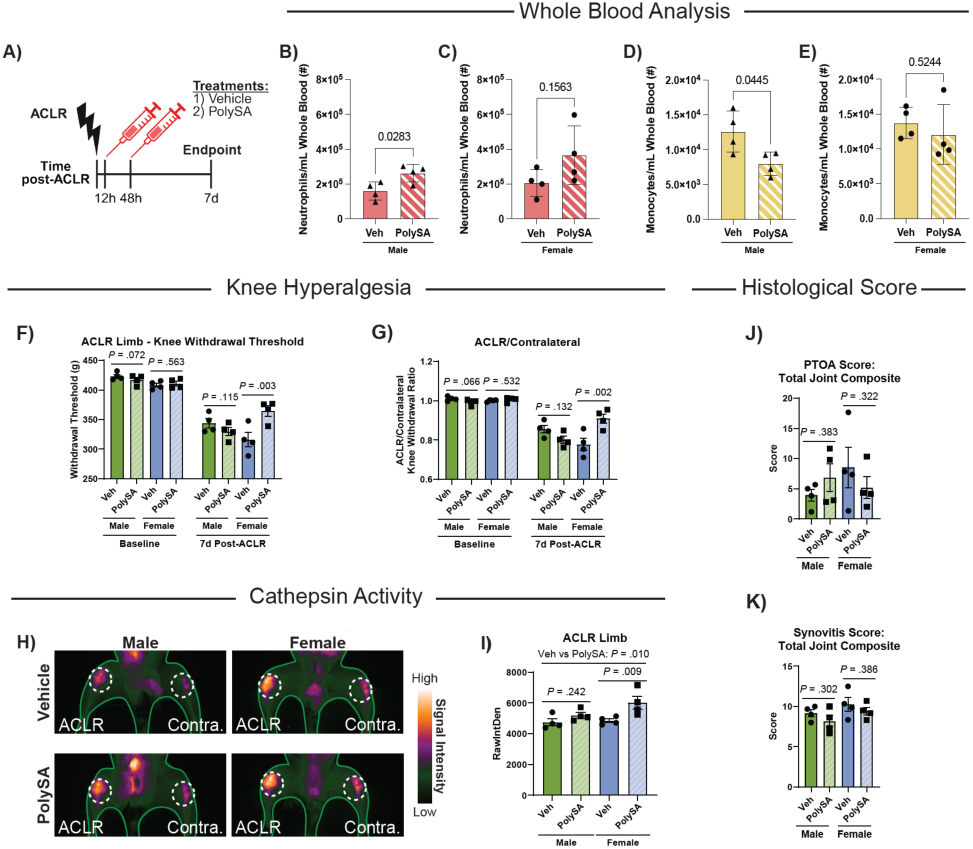
Early assessment of therapeutic efficacy of PolySA treatment. (A) Experimental schematic indicating the timing of two PolySA or Vehicle injections following ACLR, with endpoint harvests performed at 7d post-ACLR. (B-E) Whole blood neutrophil counts for (B) males and (C) females at 7d post-ACLR (n=4 per sex/treatment). Whole blood monocyte counts for (D) males and (E) females at 7d post-ACLR (n=4 per sex/treatment). Welch’s t-test run for (B-E). (F) Knee withdrawal threshold of the ACLR limb and (G) ACLR/contralateral knee withdrawal threshold ratio at baseline and 7d post-ACLR (n=4 per sex/treatment). (H) Representative images from Prosense680 NIR molecular imaging. (I) ProSense680 signal intensity quantification in the ACLR limb at 7d (n=4 per sex/treatment). Histopathologic scoring of (J) total joint PTOA severity and (K) total joint synovitis severity (n=4 per sex/treatment). Statistical analyses for F-K used linear mixed effect models. For knee withdrawal threshold, the contralateral limb was considered as a covariate for the ACLR limb and vice versa. Graphs show the mean +/- SEM.

Knee hyperalgesia testing revealed sex-dependent responses to PolySA treatment at 7d post-ACLR. Injured joints from female mice treated with PolySA exhibited significantly higher absolute withdrawal thresholds (*P*=0.003) and higher ACLR/contralateral knee withdrawal ratios (*P*=0.002), indicative of lesser injury-induced pain, relative to sex-matched vehicle-treated mice (**Fig. 2F-G**). However, this treatment effect was not observed in males (**Fig. 2F-G**). Minimal effects were observed for contralateral knee withdrawal threshold (**Suppl. Fig 3A**). PolySA treatment also had sex-specific effects on cathepsin activity, measured via ProSense680 molecular imaging. Female mice treated with PolySA exhibited ∼24% higher (*P*=0.009) ProSense signal intensity in the ACLR limb compared to the ACLR limb of sex-matched, vehicle-treated mice (**Fig. 2H-I**). This treatment effect similarly manifested in the contralateral limbs of female mice, with PolySA treated mice exhibiting ∼21% higher (*P*=0.007) contralateral limb cathepsin activity relative to sex-matched, vehicle-treated females (**Suppl. Fig. 3B**). As this increase in cathepsin is accompanied by reduced pain, it is unclear whether this effect should be deemed pro- or anti-inflammatory, and further characterization is needed to determine the phenotypic function of cathepsin activity in the context of joint injury and inflammation. Male mice did not exhibit any PolySA treatment effect on cathepsin activity in either limb (**Fig. 2H-I**). The ACLR/contralateral ProSense680 ratio was similar across treatment groups for both males and females (**Suppl. Fig. 3B**). Histopathologic evaluation showed no effect of PolySA treatment on PTOA and synovitis scores for either sex in ACLR or contralateral limbs (**Fig. 2J-K**). Representative histological images are shown in **Suppl. Fig. 4** (ACLR) and **Suppl. Fig. 5** (contralateral). Individual PTOA and synovitis subscores are shown in **Suppl. Fig. 6**. Taken together, these findings demonstrate that acute PolySA treatment blocked joint trauma-induced hyperalgesia and enhanced molecular cathepsin activity in female joints in the early post-injury period.

### PolySA-treated female mice exhibit a synovial transcriptomic signature indicative of accelerated immune resolution and matrix remodeling

Our findings that systemic PolySA treatment enhanced peri-articular cathepsin activity and exerted an early analgesic effect in female mice prompted us to assess the differentially-active molecular programs underpinning this. As synovial inflammation is strongly associated with pain in OA^6^, we focused this analysis on synovial tissue. We microdissected synovium, inclusive of Hoffa’s fat pad, from female mice and performed bulk RNA-sequencing at 7d post-ACLR. As we did not observe any therapeutic effects in male mice in the previous 7d ACLR experiment, we only conducted this experiment in female mice. Comparing synovium from mice treated with either vehicle or PolySA (at 12hrs and 48hrs) as described above, we identified 148 differentially expressed genes (DEGs, Padj < 0.05) (**Fig. 3A**). All DEGs are listed in **Suppl. Table 1** and are shown by sample in **Suppl. Fig. 7A**. Pathway overrepresentation analysis using the GO: Biological Process (BP) annotation revealed PolySA-induced enrichment of pathways indicative of cartilage anabolism (e.g. “Chondrocyte proliferation”, “Cartilage condensation”) and matrix remodeling (ex. “Extracellular matrix organization”, “Regulation of collagen biosynthetic process”) (**Fig. 3B**). PolySA-treated mice also exhibited upregulated pathways related to wound healing (e.g. “Regulation of vascular wound healing”) and susceptibility to T cell and NK cell cytotoxicity (**Fig. 3B**). Affirming successful targeting of neutrophils, PolySA-treated mice exhibited downregulation of “Neutrophil activation involved in immune response”, “Positive regulation of neutrophil degranulation”, and “Neutrophil chemotaxis” (**Fig. 3C**). Notably, neuroimmune cell (microglial, dendritic, and astrocyte cell) activation along with both innate immune responses (“Mast cell activation”,

**Figure 3:**
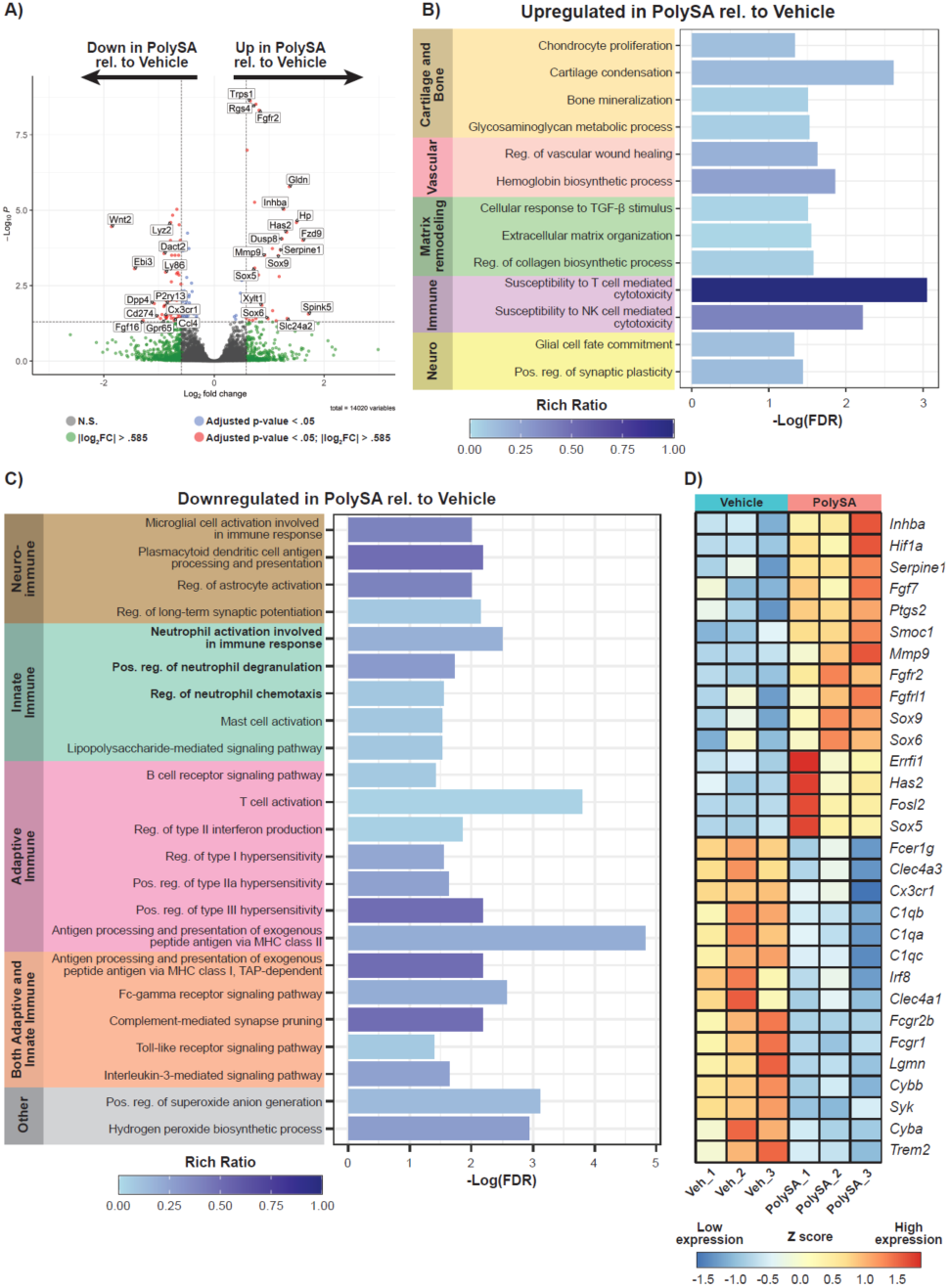
PolySA reprograms the female synovial transcriptome at 7d post-ACLR. (A) Volcano plot showing 148 differentially expressed genes (DEGs) in 7d post-ACLR synovium between Vehicle and PolySA conditions, based on bulk RNA-seq and identified using DESeq2 (adjusted p-value cutoff < 0.05). (B-C) Curated list of significant gene ontology (GO) biological process (BP) pathways associated with upregulated (B) or downregulated (C) DEGs in the PolySA group relative to Vehicle, derived using PantherDB overrepresentation analysis (−log(FDR) > 1.3). (D) Heatmap of relative expression of leading edge genes driving significant pathway results across PantherDB GO:BP, PantherDB GO:MF, and Metascape databases. Rows and columns in the heatmap were hierarchically clustered to determine sample and gene ordering. n=3 per treatment group.

“Lipopolysaccharide mediated signaling pathway”) and adaptive immune responses (“B cell receptor signaling pathway”, “T cell activation”, hypersensitivity regulation) that are known to occur downstream of neutrophil activation were also significantly downregulated in the PolySA group (**Fig. 3C**). Pathway analyses using PantherDB GO BP annotation were independently corroborated using the PantherDB Molecular Functions (MF) and Metascape databases (**Suppl. Fig. 7B-D**). Analysis of significant leading edge genes (Padj < 0.05) driving these pathway results demonstrates PolySA-induced upregulation of FGF signaling genes (*Fgf7, Fgfr2, Fgfrl1*) as well as chondrogenic genes (*Sox6, Sox9, Hif1a*) (**Fig 3D**). Surprisingly, *Mmp9*, the gene encoding the neutrophil-derived matrix protease MMP9 was also upregulated. Downregulated leading edge genes included multiple macrophage markers (*Cx3cr1, Trem2*), members of the complement system (*C1qa, C1qb, C1qc*), the interferon transcription factor Irf8, known to govern immune responses^27^, and genes encoding members of the Fc receptor family (*Fcer1b, Fcgr2b, Fcgr1*), which are critical in mediating acute immune responses{Citation} by acting as immunoglobulin receptors (**Fig 3D**).^28^ Taken together, these findings demonstrate that the beneficial therapeutic effects of PolySA are associated with molecular reprogramming of the synovium, characterized by enhanced matrix remodeling and inhibition of neuroimmune, innate, and adaptive pathways, potentially indicative of accelerated immune resolution.

### Acute intervention with PolySA mitigates long-term PTOA pain and structural damage in female mice

Finally, given our observations of the multifaceted therapeutic effects in PolySA-treated female mice in the early post-injury period, we sought to assess whether acute PolySA treatment would also mitigate long-term PTOA development. We employed the same acute treatment strategy involving two injections of vehicle or PolySA at 12hrs and 48hrs post-ACLR (**Fig. 4A**), with analyses performed at 28d post-ACLR, a timepoint of advanced pain and structural damage in the murine ACLR model.^26^ Strikingly, we observed sustained analgesic effects due to early PolySA treatment up to 28 days post-ACLR, with female mice treated with PolySA exhibiting significantly higher knee withdrawal thresholds of the ACLR limb (*P*=0.006) and significantly higher ACLR/contralateral knee withdrawal ratio (*P*=0.006) relative to the vehicle-treated group (**Fig. 4B-C**). Knee withdrawal thresholds of the contralateral limb were similar between groups (**Suppl. Fig. 8A**). Similar to our observations at 7d, no significant differences in synovial immune cell content or immune cell counts in whole blood were observed between treatment groups (**Suppl. Fig. 9-10**). We also did not observe a difference in cathepsin activity based on molecular imaging between groups in either limb or based on ACLR/contralateral signal ratio, and overall cathepsin activity was low at 28d compared to the same measurements at 7d timepoint (**Fig. 4D-E, Suppl. Fig. 8B**).

**Figure 4:**
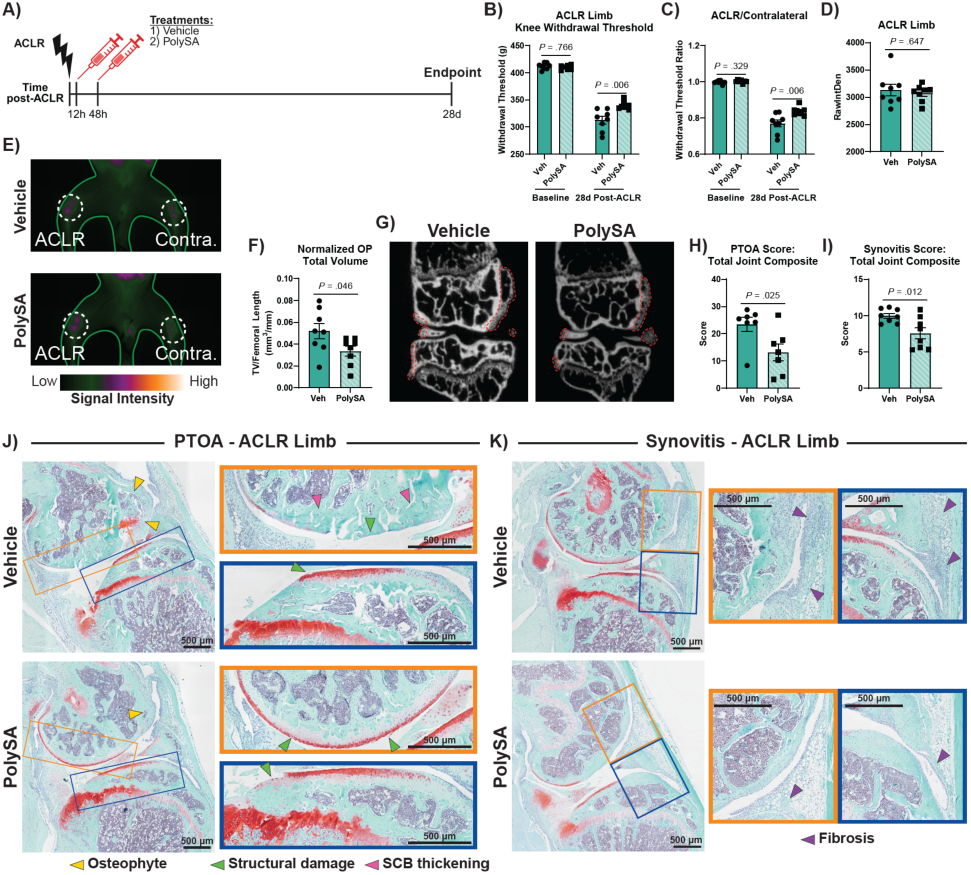
Late-stage assessment of therapeutic efficacy of PolySA treatment in females. (A) Experimental schematic indicating the timing of two PolySA or Vehicle injections following ACLR, with endpoint harvests performed at 28d post-ACLR. (B-C) Knee withdrawal threshold of the ACLR limb (B) and ACLR/contralateral knee withdrawal threshold ratio (C) at baseline and 28d post-ACLR (n=8 per treatment). (D) Signal intensity quantification from ProSense680 NIR molecular imaging in the ACLR limb at 28d (n=8 per treatment). (E) Representative images of Prosense680 signal intensity. (F-G) Normalized osteophyte volume quantification (n=8 per treatment) and (G) representative μCT images. Dashed red lines outline osteophytes. (H-I) Histopathologic scoring of total joint PTOA severity (H) and total joint synovitis severity (I) (n=7-8 per treatment). (J-K) Representative sagittal histological images for PTOA score (J) and synovitis score (K) of ACLR joints in Vehicle (top) and PolySA (bottom). Yellow arrows - osteophyte, green arrows - structural damage, pink arrows - subchondral bone thickening, purple arrows - fibrosis. Scale bar = 500μm. Statistical analyses for B-I used linear mixed effect models. For knee withdrawal threshold, the contralateral limb was considered as a covariate for the ACLR limb and vice versa. Graphs show the mean +/- SEM.

Relevant to the development of structural disease, PolySA-treated mice exhibited a marked ∼35% reduction in normalized total osteophyte volume (total volume normalized for skeletal size), with reduced osteophyte development notably observed in the medial compartment in which structural disease is maximal in this model (**Fig 4F-G**). Osteophyte maturity, measured by osteophyte bone volume fraction and mineral density, was similar between groups (**Suppl. Fig. 8C**). Histopathologic analysis of the ACLR limb revealed that PolySA-treated mice also had markedly lower total-joint PTOA scores (10.31-point difference on a max 40-point scale, *P* =.025) and synovitis scores (2.42-point difference on a max 22-point scale, *P* =.012) (**Fig. 4H-I),** evidencing a high degree of protection from structural disease. The lower total joint PTOA scores in PolySA-treated mice were primarily driven by lower articular structural damage and subchondral bone thickness of the femur and tibia in addition to decreased femoral osteophyte size (**Fig. 4J**, **Suppl. Fig. 11A**). Representative histological images demonstrate reduced articular cartilage erosion, most notably on the femoral condyles, and markedly reduced osteophyte size (**Fig. 4J**). Lower total synovitis scores in the PolySA-treated group were driven by decreased synovial fibrosis and synovial exudate (**Fig. 4K**, **Suppl. Fig. 11A**). Histological images of vehicle-treated mice show extensive synovial lining hyperplasia and fibrotic tissue in the subsynovium and infrapatellar fat pad, whereas PolySA-treated mice had a thinner lining, less subsynovial expansion, and well-preserved adipose tissue (**Fig. 4K**). As structural disease does not develop in contralateral limbs in this model^26^, no difference in total joint PTOA or synovitis scores were observed in contralateral joints between groups (**Suppl. Fig. 8D-E, 11B**). All individual PTOA and synovitis subscores are detailed in **Suppl. Fig. 11**. These findings demonstrate that early PolySA treatment following joint injury imparts sustained analgesic effects while markedly mitigating the development of structural PTOA disease.

Taken together, our overall results demonstrate that acute systemic treatment with PolySA particles targets circulating neutrophils and monocytes to reprogram the endogenous joint trauma response, resulting in the mitigation of pain, altering pathological synovial gene expression, and ameliorating the development of structural disease.

## Discussion

Using a murine model of joint injury, we showed that acute intervention with systemically-administered PolySA particles can prevent the development of PTOA-related pain, synovitis, and structural degeneration in a sex-specific manner. We observed that only female mice responded to the PolySA particle therapeutic, where a divergence in treatment effect occurred by 7d post-ACLR. This sex-specific response was particularly surprising considering that treatment with PolySA at 12hrs post-ACLR resulted in similar synovial neutrophil, monocyte, and macrophage counts at 24hrs post-ACLR for males and females. Prior studies utilizing PolySA particles in acute lung injury and thrombosis models have not examined potential sex-specific effects of neutrophil activation and recruitment in response to PolySA^19,21–23^, highlighting the importance of this study for the clinical translation of PolySA particle-based therapeutics. While our study showed no sex-based effect of PolySA on neutrophil recruitment to the synovium or circulating neutrophil counts, a study of ACLR followed by ACL reconstruction showed sex-dependent effects of acute adenosine, lidocaine, and magnesium (ALM) therapy on circulating immune cells.^29^ Observing no difference in our study of synovial immune cell counts at 24hrs post-ACLR between PolySA treated males and females, the therapeutic effects observed in females at 7d post-ACLR (increased cathepsin activity, alleviation of pain) are likely due to sex-specific, PolySA-dependent molecular reprogramming of female systemic immune cells and synovial cells. The impact of sex on acute and chronic inflammatory conditions has been well characterized, where human neutrophils exhibit distinct sex-specific inflammatory phenotypes, with female neutrophils incurring a more robust response to inflammation.^30,31^ This is evidenced in the higher prevalence of autoimmune disorders and inflammatory conditions in females, where human females experience worse PTOA prognoses.^32,33^

Cathepsin activity as measured by ProSense680 was increased in female mice that received PolySA treatment as compared to vehicle treated female mice at 7d (**Fig. 2H-I**, **Suppl. Fig. 3B**). This was a particularly surprising finding as our bulk RNA seq data at 7d shows the significant (adjusted p-value <.05) downregulation of cathepsin C (*Ctsc*, Log_2_FC = -0.464) and cathepsin S expression (*Ctss*, Log_2_FC = -0.516) in the PolySA treated group relative to vehicle (**Suppl. Table 1**), which indicates that while cathepsin family transcript expression does not increase, the activity of the already present cathepsin enzymes was increased with PolySA treatment. Cathepsin C (*Ctsc*), also known as dipeptidyl peptidase 1 (*Dpp1*) is responsible for the cleavage and activation of neutrophil serine proteases such as cathepsin G.^34^ Cathepsin G has been shown to degrade lubricin (PRG4), a key protectant against synovial joint disease, derived from the synovial fluid of OA patients *in vitro*.^35^ Using a mouse model of myelin oligodendrocyte protein-induced neuropathic pain, Harada, et. al., demonstrated that cathepsin E knockout mice were protected against mechanical allodynia.^36^ Furthermore, they demonstrated that specifically neutrophil-derived cathepsin E mediated this pain sensitization and this is in part due to the requirement for cathepsin E in neutrophil elastase enzymatic activity.^36^ Analysis of human synovial fluid and serum from non-OA cadavers, early- and late-stage OA patients, and rheumatoid arthritis (RA) patients showed elevation of cathepsin B and cathepsin S in both OA and RA patients with cathepsin S showing greater correlation expression in RA than OA patients.^37^ Furthermore, cathepsin K has been shown to play a role in the progression of OA, acting as a collagenase and aggrecan-cleaving protease, which contributes to the degradation of articular cartilage.^38^ Thus, further exploration of which cell-type specific cathepsins are changing their activity at 7d post-ACLR is necessary to determine how increased cathepsin activity due to PolySA treatment manifests in synovitis and PTOA pathophysiology.

Our prior work has identified a difference in male and female knee hyperalgesia in this model of non-invasive ACLR, where male mice exhibited significantly lower withdrawal thresholds than female mice in ACLR limbs at 28d post-ACLR.^26^ These results were attributed to distinct differences in the synovial transcriptome, where female mice had a more pronounced resolution of synovial inflammation. Histopathological scoring at 7d post-injury indicated no difference in synovitis progression for female mice that could help account for differences in hyperalgesia between the PolySA and vehicle groups. Further, total immune cell counts within the joint showed no impact of treatment. However, this flow cytometric analysis only characterized major cell markers to identify immune cell populations and did not measure the precise phenotype of these immune cells. Thus, bulk RNAseq was used to further characterize the inflammatory environment for vehicle treated female mice as compared to those treated with PolySA. Bulk RNAseq identified indicators of synovial immune response resolution (downregulation of *Cx3cr1, Lyz2, Cd4*) for PolySA treated females that gave insight to the altered progression of inflammation. Further, the downregulation of innate and adaptive immune response pathways, which occur downstream of neutrophil activation, support that PolySA particle treatment successfully targets neutrophils and alters their interaction with the synovial cell niche. Lastly, differences in whole blood cell counts for males versus females showed that males maintained significantly altered neutrophil and monocyte counts for PolySA versus vehicle treated mice, where females appeared to reach baseline counts by 7d post-injury. Future studies should further characterize the specific mechanisms by which PolySA has an effect on immune cells in female mice, and determine a means to produce similar results in male mice, thereby facilitating the use of PolySA in the clinic.

Pain associated with osteoarthritis has also been attributed to immune cell presence, namely macrophages, within the dorsal root ganglia (DRG).^39,40^ However, neutrophils have also been reported to contribute DRG activation as Harada, et. al., also demonstrated that neutrophil accumulation in the DRG is sufficient to induce mechanical allodynia in mice.^36^ While the direct impact of neutrophils in the DRG on pain progression in PTOA is not well characterized, other works have identified neutrophils as contributors to chronic pain.^41^ Employing a systemic PolySA treatment in our studies rather than a local intra-articular injection emphasizes the need for future experiments investigating the role of PolySA-mediated neutrophil activation in the DRG and how this contributes to alleviation of pain with PolySA treatment in PTOA.

In conclusion, our findings demonstrated that PolySA successfully alters neutrophil-driven disease mechanisms in PTOA to provide sustained pain alleviation mitigation of structural damage in a sex-specific manner. Thus, targeting immune cells prior to their infiltration into injured joints during the acute phase of inflammation following trauma represents a promising approach for the prevention of PTOA.

## Methods

### Human study approvals

Human blood was collected from healthy male and female donors 20-30 years old under a protocol approved by the University of Michigan Internal Review Board (IRB: #HUM00013973). Informed and written consent was collected from each donor prior to donation. All donors were compensated monetarily for their participation.

### Fabrication of Poly(Salicylic Acid) microparticles

PolySA polymer was fabricated by the Uhrich group at UC Riverside. PolySA microparticles were fabricated using an oil in water emulsion, where PolySA was initially dissolved into dichloromethane (DCM) and then slowly added to the emulsified water phase (1% polyvinyl alcohol). The emulsion was mixed for 2 hours to facilitate evaporation of DCM, and then particles were collected and cleaned via subsequent centrifugation steps in deionized water. All particles were passed through a 2 µm filter prior to lyophilization. Cy5.5 labeled particles were fabricated in an equivalent manner, with the addition of 1% Cy5.5 conjugated PolySA polymer to DCM/polymer mixture. PolySA polymer was conjugated with Cy5.5 amine (Lumiprobe, Catalog # 470C0) using carbodiimide crosslinker chemistry, consistent with previously described methods.^22^

### Characterization of PolySA microparticles

Physical characterization of PolySA particles was completed using JEOL JSM-7800FLV scanning electron microscope (SEM) and a Malvern Zetasizer. Particle sizing was measured using ImageJ, where >100 particle diameters were measured for both PolySA and Cy5.5-PolySA.

### PolySA treatment of activated human monocytes

Whole blood was collected from healthy human donors in EDTA anticoagulant tubes (BD, Catalog # 366643) and placed into a 37°C incubator for 1 hour. Following incubation, whole blood was centrifuged at 200g for 15 minutes to collect the buffy coat containing peripheral blood mononuclear cells and placed into a 5 mL round bottom tube.

A monocyte isolation kit (Stemcell Technologies, Catalog # 19669) was used, where 110 µL antibody solution and 50 µL magnetic beads were incubated with blood for 5 minutes. The whole blood suspension was then diluted with 1 µM EDTA in phosphate buffered saline and placed into a magnetic separator for 3 minutes. Supernatant was poured into a new 5 mL round bottom tube, incubated with an additional 50 µL magnetic beads for 5 minutes, and then placed into a magnetic separator for 3 minutes. Supernatant was poured into a new 5 mL round bottom tube and then placed into a magnetic separator for 3 minutes, where remaining supernatant composed isolated monocytes. Monocytes were centrifuged, washed, and resuspended in RPMI supplemented with 10% donor plasma at a concentration of 1x10^6^ cells/mL. 100,000 cells were placed into a microcentrifuge tube corresponding to each experimental group. Unactivated monocytes received treatment only. Activated monocytes received 5000 pg/mL TNF-αalongside treatment. Treatments included PolySA particles at a particle:cell ratio of 16:1, free salicylic acid at a concentration of 0.05 mg/mL (equivalent to total salicylic acid contained in 1.6x10^6^ PolySA particles), and 1 µm polystyrene particles at a particle:cell ratio of 16:1. Treatments and TNF-⍺ were administered to cells simultaneously and then cells were incubated for 1 hour at 37°C and 5% CO_2_ prior to staining for flow cytometry. 5 donors were used for each experimental group.

### Flow cytometry of activated human monocytes

Following treatment, isolated human monocytes were stained for flow cytometric analysis. Briefly, cells were treated with human FcX and then stained using the antibody panel shown in **Suppl. Table 2**. Samples were fixed using 1-Step Fix/Lyse (eBioscience, Catalog # 00-5333-54), washed, and resuspended in FACS buffer (PBS (-/-) + 2% FBS). Samples were analyzed using an Attune flow cytometer and further analysis was conducted using FlowJo to determine particle positive populations and overall relative surface protein expression as compared to vehicle treated, non-activated samples. Sample gating strategy shown in **Suppl. Fig. 12**.

### Mice

All animal studies were conducted with approval of the University of Michigan Institutional Animal Care and Use Committee. Mice were housed in a 12 hr light/dark facility with ad libitum access to food and water. Male and female C57/Bl6J mice (Jackson Laboratories) aged 12-14 wks old were used for all studies. Mice were randomized to treatment groups within each experiment.

### Noninvasive anterior cruciate ligament rupture (ACLR)

Noninvasive anterior cruciate ligament rupture (ACLR) was used to induce PTOA as described previously.^42^ Briefly, mice were anaesthetized (5% isoflurane induction, 2% isoflurane maintenance) and placed prone on a custom fixture. The hindlimb receiving ACLR was fixed at 100° of flexion and subject to a pre-conditioning cycle (CellScale Univert S2, CellScale, Waterloo, ON, Canada). Following pre-conditioning, a single mechanical overloading event caused anterior tibial subluxation and subsequent ACL rupture. Unilateral ACLR (uniACLR) mice received carprofen (5 mg/kg) and bilateral ACLR (biACLR) mice received buprenorphine (3.25 mg/kg). Sham mice received anaesthesia and analgesia only, no loading cycle. At the endpoint, mice were euthanized via CO_2_ asphyxiation. All experimental mouse cohorts are summarized in **Table 1**.

**Table 1.**
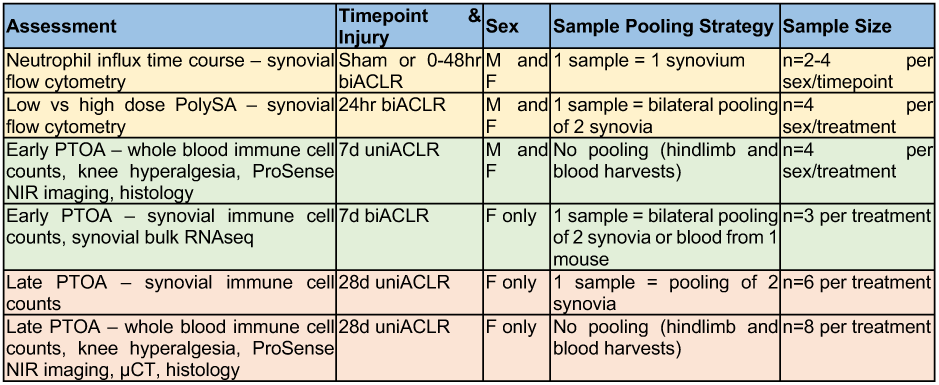
Methods summary by experimental mouse cohort. For each experiment, type of assessment, timepoint, injury, sex, sample pooling, and sample size are specified. M – male. F – female.

| Assessment | Timepoint & Injury | Sex | Sample Pooling Strategy | Sample Size |
| --- | --- | --- | --- | --- |
| Neutrophil influx time course – synovial flow cytometry | Sham or 0-48hr biACLR | M and F | 1 sample = 1 synovium | n=2-4 per sex/timepoint |
| Low vs high dose PolySA – synovial flow cytometry | 24hr biACLR | M and F | 1 sample = bilateral pooling of 2 synovia | n=4 per sex/treatment |
| Early PTOA – whole blood immune cell counts, knee hyperalgesia, ProSense NIR imaging, histology | 7d uniACLR | M and F | No pooling (hindlimb and blood harvests) | n=4 per sex/treatment |
| Early PTOA – synovial immune cell counts, synovial bulk RNAseq | 7d biACLR | F only | 1 sample = bilateral pooling of 2 synovia or blood from 1 mouse | n=3 per treatment |
| Late PTOA – synovial immune cell counts | 28d uniACLR | F only | 1 sample = pooling of 2 synovia | n=6 per treatment |
| Late PTOA – whole blood immune cell counts, knee hyperalgesia, ProSense NIR imaging, μCT, histology | 28d uniACLR | F only | No pooling (hindlimb and blood harvests) | n=8 per treatment |

### Synovium digestion and flow cytometry

Synovia were harvested 0-48 hours post-ACLR, digested, stained, and immune cell content was quantified by flow cytometry. A separate cohort of mice were administered either vehicle, low dose PolySA, or high dose PolySA treatment at 12 hrs post-ACLR and synovia and whole blood were harvested 24 hrs post-ACLR for digestion, staining, and immune cell quantification by flow cytometry.Two additional cohorts of mice received vehicle or PolySA treatment at 12 hrs and 48hrs post-ACLR and synovia and whole blood were harvested 7d or 28d post-ACLR for digestion, staining, and immune cell quantification by flow cytometry.

Briefly, anterior synovium including Hoffa’s fat pad were microdissected from the knee joint. Synovia were weighed to determine total mass of collected tissue. After weighing, synovia were digested in a solution of Liberase (Millipore Sigma, Catalog # 5401127001), DNase (Millipore Sigma, Catalog # DN25), and Collagenase IV (Millipore Sigma, Catalog # C5138) for 35 minutes while stirred at 200 rpm at 37°C and mixed via gentle pipetting every 10 minutes. Once synovial tissue was digested, the cell suspension was washed with FACS buffer and incubated with FcX for 5 minutes before staining with an immune cell antibody panel (**Suppl. Table 3**). Stained samples were fixed using 1-Step Fix/Lyse, resuspended in FACS buffer, and run on an Attune flow cytometer. Total cell counts from flow cytometry were determined by recording volume of sample run relative to total cell suspension volume. Total cell counts were then divided by mass of tissue to obtain a relative cell count across joint samples. Analysis was performed in FlowJo to quantify synovial immune cell populations. Sample gating strategy is shown for non fluoro-labelled PolySA particle treatment **Suppl. Fig. 13** and fluoro-labelled PolySA particle treatments in **Suppl. Fig. 14**.

### Whole blood collection and flow cytometry

Whole blood was collected from mice via cardiac puncture using 50 µL heparin solution (0.762 mg/mL in PBS (-/-)) and stored on ice until processed for flow cytometry. 50 µL whole blood was aliquoted and treated with mouse FcX for 10 minutes prior to staining with antibody panel. Antibody panel shown in **Suppl. Table 3**. Stained samples were fixed and red blood cells were lysed using 1-Step Fix/Lyse. Samples were washed and resuspended in FACS buffer. Total cell counts from flow cytometry were determined by recording volume of sample run relative to total cell suspension volume. Analysis was performed in FlowJo to quantity neutrophil and monocyte populations in whole blood. Sample gating strategy is shown in **Suppl. Fig. 15**.

### Retro-orbital PolySA treatments

At 12 hrs post-ACLR, mice were administered either vehicle (PBS (-/-)), low dose PolySA (2x10^8^ particles/animal in 100 µL), or high dose PolySA (4x10^8^ particles/animal in 100 µL) treatment and synovia were harvested 24 hrs post-ACLR to quantify immune cell influx by flow cytometry. Separate cohorts of mice were given two acute doses of vehicle (PBS (-/-)) or PolySA (4x10^8^ particles/animal in 100 µL) retro-orbitally at 12 hrs and 48 hrs post-ACLR and then evaluated at 7d or 28d post-ACLR as summarized in **Table 1**.

### Near-infrared imaging of cathepsin activity

Twenty-four hours prior to harvest mice were retro-orbitally injected (100 µL) with ProSense 680 (Revvity, #NEV10003), a cathepsin-activated near-infrared probe. At harvest, either 7d or 28d post-ACLR, mice were euthanized, skinned, and NIR imaging (Pearl Impulse, LI-COR, Lincoln, NE, United States) was performed (7d: n=4 per sex/treatment, 28d: n=8 females per treatment). Images were analyzed by a blinded observer in ImageJ. Briefly, a consistently sized and shaped ROI was drawn over the knee and the raw integrated density (RawIntDen, RID) was recorded. The contralateral/ACLR RID ratio was calculated for each image.

### Knee hyperalgesia testing

Knee hyperalgesia testing was performed using a Randell-Selitto device (IITC Life Science, Woodland Hills, CA) with a custom blunt tip probe as described previously^26^ at baseline and either 7d (n=4 per sex/treatment) or 28d (n=8 females per treatment) post-ACLR. Briefly, consistently increasing pressure was applied to the medial aspect of the knee and when the mouse struggled or vocalized, the force, termed “knee withdrawal threshold”, was recorded. Three replicates were recorded and averaged together for each limb/timepoint. Assessments were performed by a blinded observer.

### Hindlimb collection

Injured and uninjured contralateral hindlimbs were harvested either 7d or 28d post-ACLR by disarticulating the limb at the hip, fixed for 48 hrs in 10% neutral buffered formalin while nutating at 4°C. Following fixation, limbs were rinsed in running water for 5 min and stored in 70% ethanol (EtOH) at 4°C until decalcification or micro computed tomography scanning.

### Micro-computed tomography

Limbs from 28d ACLR mice were scanned using micro-computed tomography (μCT, SkyScan 8.9 μM voxel size, n=8 females per treatment). Two hours prior to scanning, limbs were transferred from 70% EtOH into scanning cassettes in PBS. Scanning was performed and scans were reconstructed. Limbs were transferred back into 70% EtOH at 4°C until ready for decalcification. Osteophytes (OP) were manually outlined by a blinded evaluator, and the femoral length was measured using Dragonfly. Normalized osteophyte volume was calculated by dividing the total OP volume by femoral length to account for differences in skeletal size.

### Paraffin histology and histopathological scoring

Limbs from 7d (n=4 per sex/treatment) and 28d (n=7-8 females per treatment) mice were decalcified in 10% ethylene diamine tetraacetic acid (EDTA) while nutating at 4°C for 2 weeks with a solution refresh at 1 week. Samples were washed under running water for 5 min and stored in 70% EtOH at 4°C until ready for paraffin processing. Limbs were paraffin processed at the Michigan Integrative Musculoskeletal Health Core Center’s (MiMHC) Structure, Composition, and Histology Core (NIAMS P30 AR069620). Following processing, limbs were embedded in paraffin, sectioned sagittally (5 μm thickness), stained with Safranin-Orange/Fast Green, and imaged (n=2-8 sections per limb spanning the medial condyle, Eclipse Ni E800 with DS-Ri2 camera, Nikon, Tokyo, Japan). Histological assessment of PTOA (**Suppl. Table 4**) and synovitis severity (**Suppl. Table 5**) was performed by a blinded observer using scoring systems based on previous studies.^26,43^ All histopathological scoring system expansions and criterion specification were determined by LL.

### Flow sorting of live cells and RNA isolation for bulk RNA sequencing

Mice were treated at 12 hrs and 48 hrs post-ACLR with vehicle or PolySA and at 7d post-ACLR synovia were harvested for flow cytometry cells and bulk RNA sequencing. Dissected synovia were digested and stained (**Suppl. Table 3**) for flow cytometry to assess immune cell populations as described above. Sample gating strategy is shown in **Suppl. Fig. 14**. Live cells were sorted into ice cold RPMI supplemented with 10% FBS. Samples were kept on ice until centrifuged to pellet cells, and then supernatant was removed. Pelleted cells were resuspended and homogenized in RLT buffer (info) and stored at -80°C until full RNA isolation was completed using the Qiagen RNeasy isolation kit (Qiagen, Catalog # 74004). Isolated RNA was then stored at -80°C until processed for bulk RNA sequencing.

### Bulk RNA sequencing and bioinformatics analysis

Total RNA was extracted from all samples and assessed for quality using RNA integrity number (RIN) values, with only samples exhibiting RIN ≥ 8 included in subsequent analyses. Genes with counts ≥ 10 in at least 3 samples were retained for downstream analysis. Outliers were assessed via principal component analysis (PCA), and significant statistical outliers were not detected (**Suppl. Fig. 16A**). The top ten top and bottom gene loadings for PC1 and PC2 are shown (**Suppl. Fig. 16B**). To assess potential contamination from surrounding muscle tissue and evaluate sample heterogeneity, a muscle enrichment score was calculated based on a predefined list of muscle-enriched genes (**Suppl. Fig. 16C**). Sample correlation was visualized using a Manhattan distance matrix (*pcaExplorer* v3.2.0, **Suppl. Fig. 16D**). DEGs between treatment groups (PolySA vs. Vehicle) were calculated using *DESeq2* (v1.48.1) with default parameters. DEGs were defined as those with an adjusted p-value < 0.05. Pathway enrichment analysis was conducted using the PantherDB statistical overrepresentation test (GO biological process complete (BP), GO molecular functions (MF)), or Metascape, with either upregulated (Log_2_FC > 0) or downregulated (Log_2_FC < 0) genes (*P_adj_* < 0.05) as input. Gene set enrichment analysis (GSEA) was conducted (*clusterProfiler* v4.16.0) using all genes, ordered by a calculated Log_2_-Fold Change (calculated from *DESeq2*), minimum gene set size of 10, maximum gene set size of 800, and p-value cutoff < 0.05. All visualizations, including volcano plots, heatmaps, and other figures, were generated using the *EnhancedVolcano* (v1.26.0), *ggplot2* (v4.0.0), and *pheatmap* (v1.0.13) packages in R.

### Statistical analysis

SPSS (v29, IBM, Armonk, NY, United States) and Prism 10.0 (Graphpad, San Diego, CA) were used for statistical analyses. All data points represent independent biological replicates for both human and mouse studies. Analysis of immune cell infiltration into synovial tissue and whole blood cell counts for 24 hour studies were completed using a One-Way ANOVA with post hoc Fisher’s LSD. All other comparisons for immune cell infiltration into tissue and whole blood cell counts were conducted using Welch’s t-test. Statistical analysis of isolated monocyte activation was conducted using a One-Way ANOVA with post hoc Tukey’s test. Linear mixed effect models within cohort (7d, 28d) were used to compare knee hyperalgesia, NIR imaging, histopathological scoring, and μCT data between treatment groups. For knee withdrawal threshold, the contralateral limb was considered as a covariate for the ACLR limb and vice versa.

## Funding

National Institutes of Health NIAMS (R21 AR080502)

National Science Foundation Graduate Research Fellowship grant (DGE 2241144) (LL)

National Institutes of Health T32 (T32 GM145304) (MLF)

Michigan Integrative Musculoskeletal Health Core Center’s (MiMHC) Structure, Composition, and Histology Core (NIAMS P30 AR069620)

University of Michigan, Department of Orthopaedic Surgery

## Author contributions

Conceptualization: MLF, LL, OEA, TM

Methodology: MLF, LL, OEA, TM

Investigation: MLF, LL, AAG, MVG, SCH, IJS

Visualization: MLF, LL, AAG

Funding acquisition: MLF, LL, OEA, TM

Writing - original draft: MLF, LL

Writing - review and editing: MLF, LL, OEA, TM

## Data and materials availability

Bulk RNA-seq data will be publicly available on the National Institutes of Health Gene Expression Omnibus (GEO) database repository.

