## Supplemental File for "Acute immune modulation with poly-salicylic acid particles ameliorates pain and structural damage in post-traumatic osteoarthritis"

### Supplemental Figures

A)

| Particle Type | Diameter (µm) | Zeta Potential (mV) |
|---|---|---|
| PolySA | 1.14 ± 0.33 | -16.5 |
| PolySA-Cy5.5 | 1.18 ± 0.28 | -21.0 |

B)

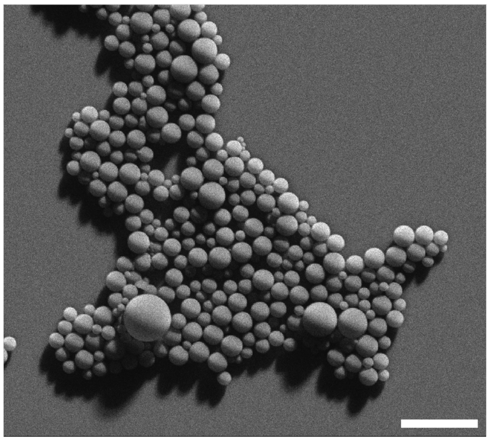

**Suppl. Fig. 1. PolySA particle characterization.** (A) Particle size and zeta potential distribution. SEM image of (B) Cy5.5-PolySA particles. Scale bar is 5 µm.

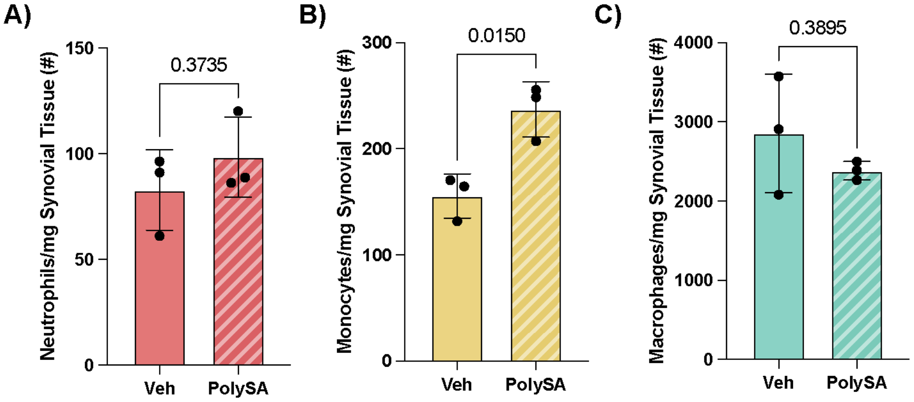


**Suppl. Fig. 2. Synovial immune cell influx 7d post-ACLR.** Counts are shown for (A) neutrophils, (B) monocytes, and (C) macrophages in the synovium at 7d post-ACLR with vehicle or PolySA treatments occurring at 12hrs and 48hrs post-ACLR for female mice.

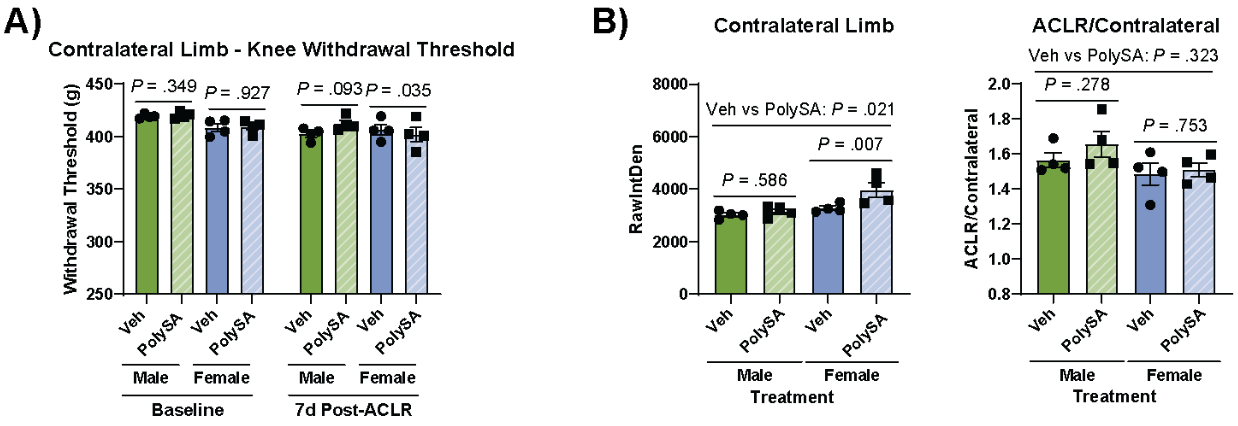


**Suppl. Fig. 3. 7d post-ACLR *in vivo* phenotyping supplement. (**A) Knee withdrawal threshold of the contralateral limb at baseline and 7d post-ACLR (n=4 per sex/treatment). (B) ProSense680 signal intensity quantification in the contralateral limb and the ACLR/contralateral ProSense680 signal ratio at 7d (n=4 per sex/treatment). Graphs show the mean +/- SEM.

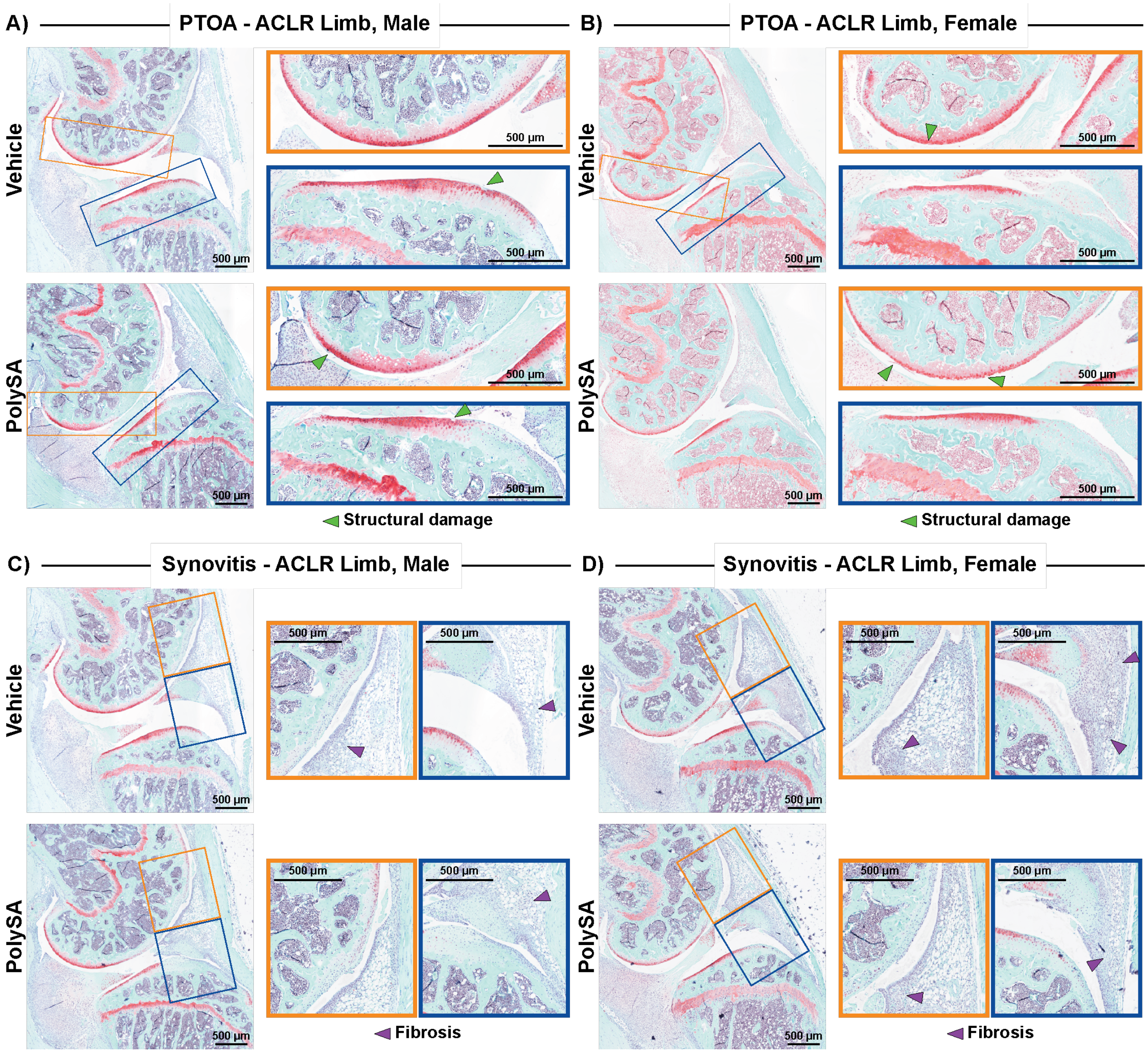


**Suppl. Fig. 4. 7d post-ACLR representative histology images of the ACLR limb.** Representative images depict an ACLR limb with a (A-B) PTOA severity score or (C-D) synovitis score closest to the treatment group mean. Green arrows - structural damage, purple arrows - fibrosis. Scale bar = 500µm.

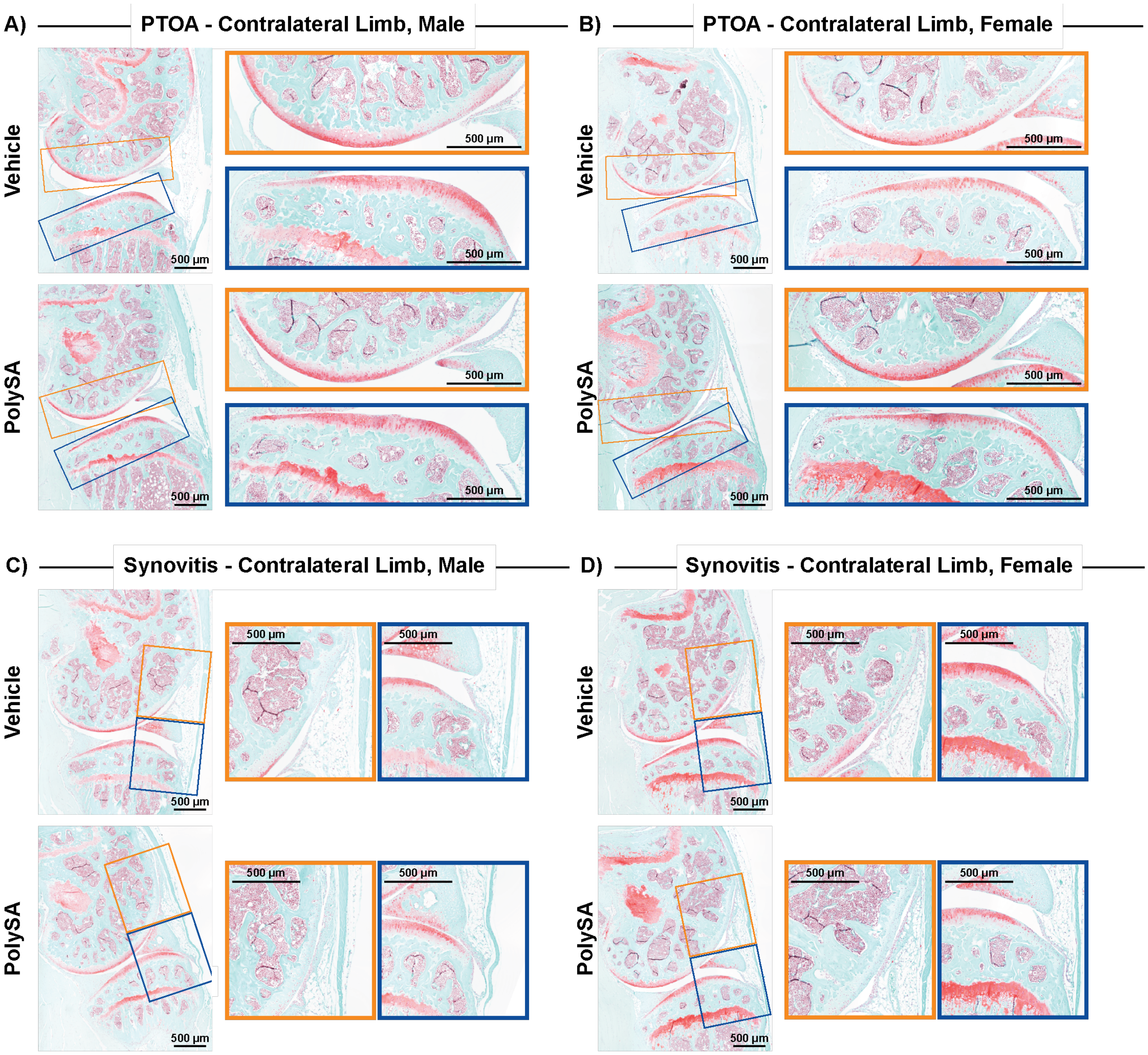


**Suppl. Fig. 5. 7d post-ACLR representative histology images of the contralateral limb.** Representative images depict a contralateral limb with a (A-B) PTOA severity score or (C-D) synovitis score closest to the treatment group mean. Scale bar = 500µm.

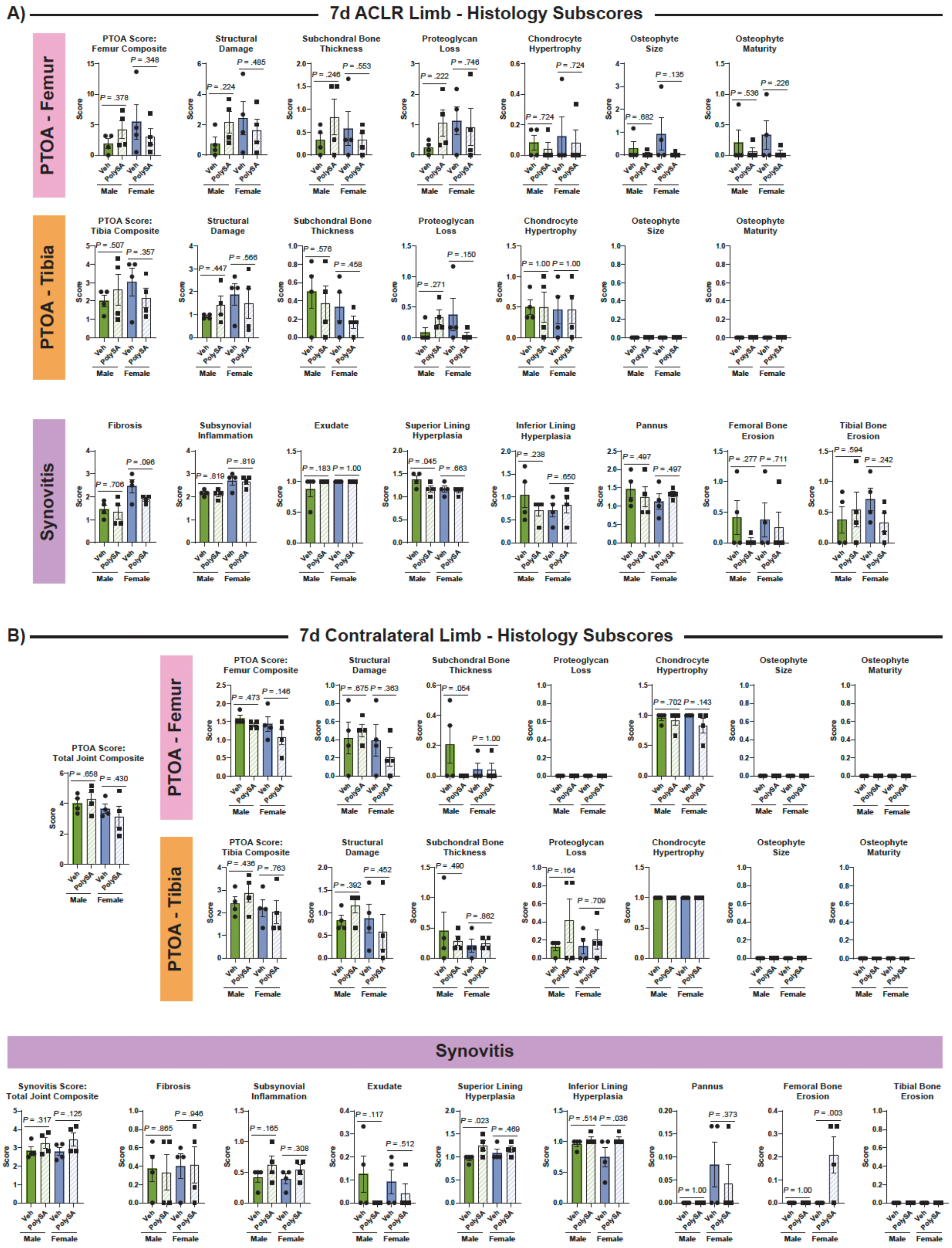


**Suppl. Fig. 6. 7d post-ACLR histopathology subscores.** Graphs show each individual subscore of PTOA and synovitis severity at 7d post-ACLR for the (A) ACLR limb and the (B) contralateral limb (n=4 per sex/treatment). Graphs show the mean +/- SEM.

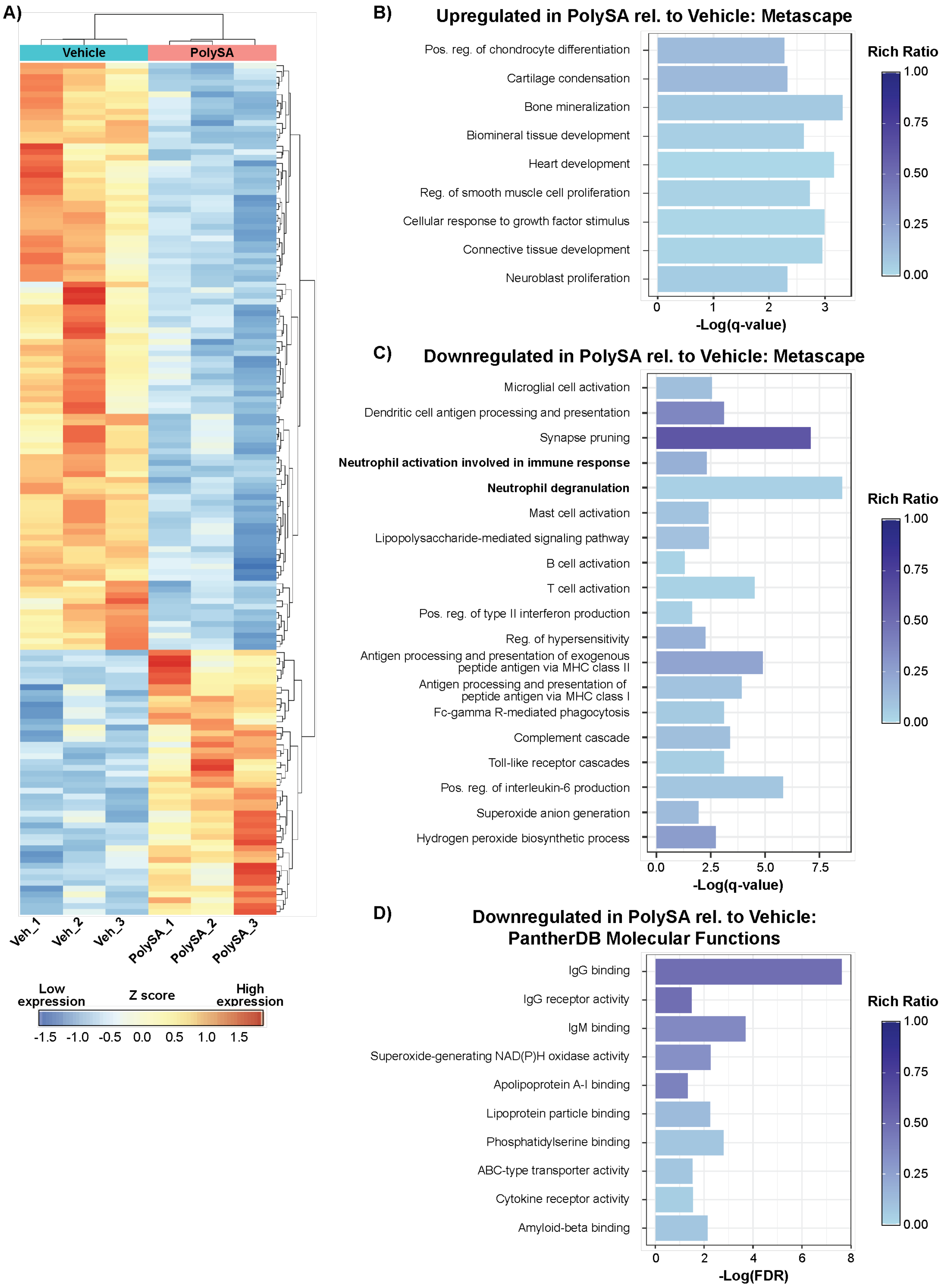

A)
Vehicle
PolySA
Veh_1
Veh_2
Veh_3
PolySA_1
PolySA_2
PolySA_3
Low expression
Z score
High expression
-1.5 -1.0 -0.5 0.0 0.5 1.0 1.5
B)
Upregulated in PolySA rel. to Vehicle: Metascape
Pos. reg. of chondrocyte differentiation
Cartilage condensation
Bone mineralization
Biomineral tissue development
Heart development
Reg. of smooth muscle cell proliferation
Cellular response to growth factor stimulus
Connective tissue development
Neuroblast proliferation
0 1 2 3
-Log(q-value)
Rich Ratio
1.00
0.75
0.50
0.25
0.00
C)
Downregulated in PolySA rel. to Vehicle: Metascape
Microglial cell activation
Dendritic cell antigen processing and presentation
Synapse pruning
Neutrophil activation involved in immune response
Neutrophil degranulation
Mast cell activation
Lipopolysaccharide-mediated signaling pathway
B cell activation
T cell activation
Pos. reg. of type II interferon production
Reg. of hypersensitivity
Antigen processing and presentation of exogenous peptide antigen via MHC class II
Antigen processing and presentation of peptide antigen via MHC class I
Fc-gamma R-mediated phagocytosis
Complement cascade
Toll-like receptor cascades
Pos. reg. of interleukin-6 production
Superoxide anion generation
Hydrogen peroxide biosynthetic process
0.0 2.5 5.0 7.5
-Log(q-value)
Rich Ratio
1.00
0.75
0.50
0.25
0.00
D)
Downregulated in PolySA rel. to Vehicle: PantherDB Molecular Functions
IgG binding
IgG receptor activity
IgM binding
Superoxide-generating NAD(P)H oxidase activity
Apolipoprotein A-I binding
Lipoprotein particle binding
Phosphatidylserine binding
ABC-type transporter activity
Cytokine receptor activity
Amyloid-beta binding
0 2 4 6 8
-Log(FDR)
Rich Ratio
1.00
0.75
0.50
0.25
0.00

**Suppl. Fig. 7. 7d bulk RNAseq biological analysis.** (A) Heatmap showing expression of all differentially expressed genes (DEGs) identified between PolySA vs. Vehicle in 7d post-ACLR synovium (p-adjusted < 0.05). (B-C) Selected Metascape pathways for both upregulated ($Log_2FC > 0$) (B) and downregulated ($Log_2FC < 0$) (C) significant terms (-log(q-value) > 1.3). (D) Significant (-log(FDR) > 1.3) PantherDB gene ontology (GO) molecular function (MF) pathways for downregulated ($Log_2FC < 0$) DEGs. Only one significant upregulated ($Log_2FC > 0$) pathway was uncovered in this analysis (protein binding) and is not shown.

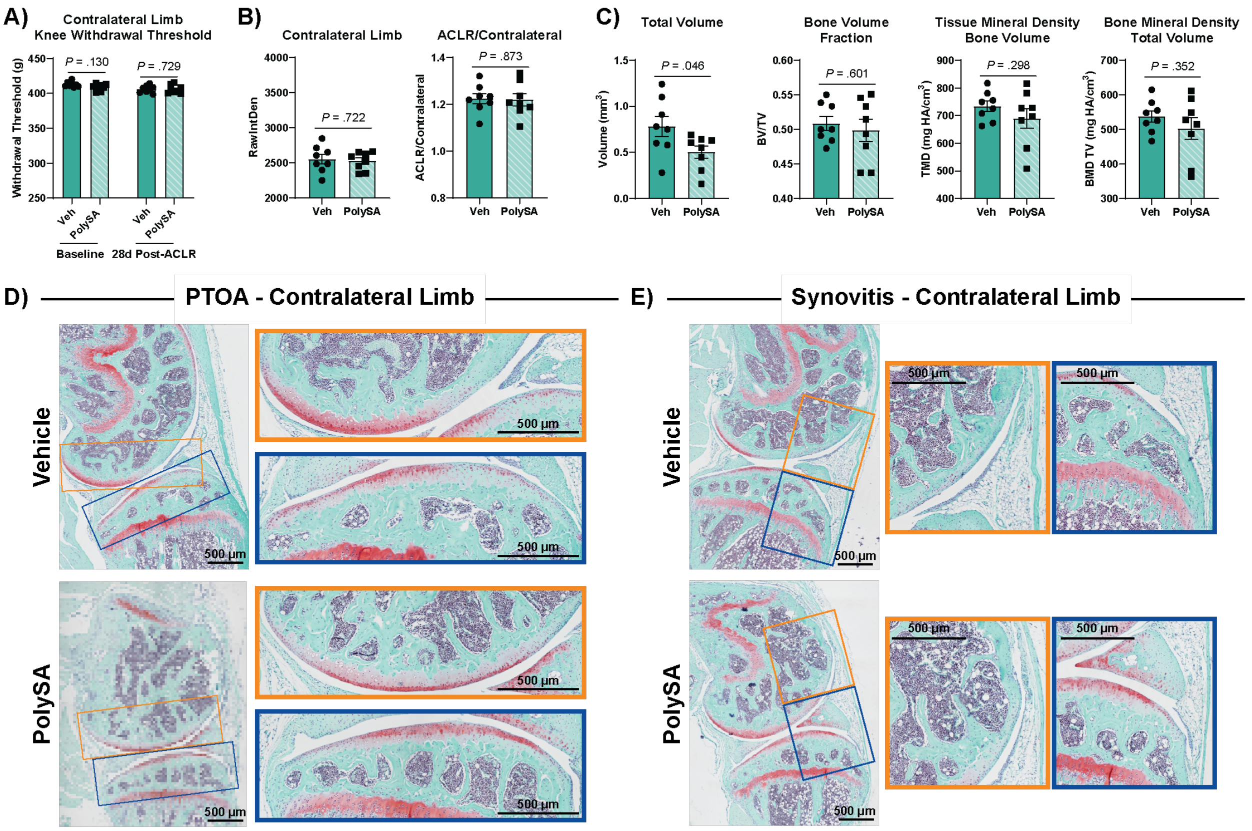


**Suppl. Fig. 8. 28d post-ACLR phenotyping supplement.** (A) Knee withdrawal threshold of the contralateral limb at baseline and 28d post-ACLR (n=8 per treatment). (B) ProSense680 signal intensity quantification in the contralateral limb and the ACLR/contralateral ProSense680 signal ratio at 28d (n=8 per treatment). (C) Additional osteophyte quantification parameters (n=8 per treatment). Representative images depict a contralateral limb with a (D) PTOA severity score or (E) synovitis score closest to the treatment group mean. Scale bar = 500µm. Graphs show the mean +/- SEM.

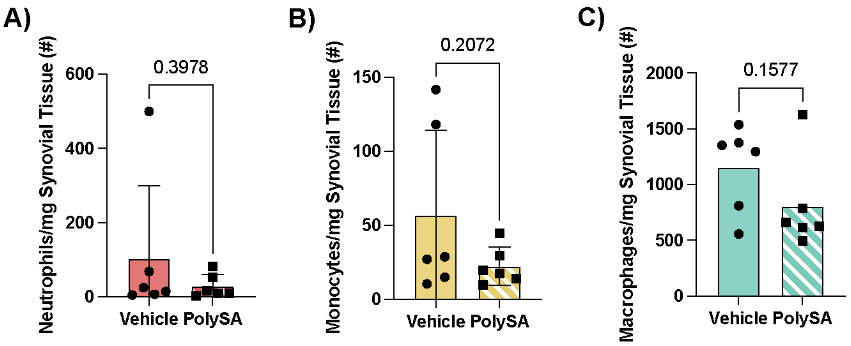


**Suppl. Fig. 9. Synovial immune cell influx 28d post-ACLR.** Counts are shown for (A) neutrophils, (B) monocytes, and (C) macrophages in the synovium at 28d post-ACLR with vehicle or PolySA treatments occurring at 12hrs and 48hrs post-ACLR.

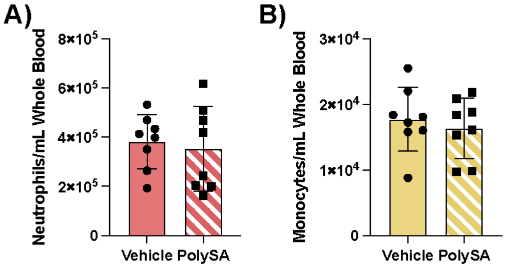


**Suppl. Fig. 10. Neutrophil and monocyte cell counts in whole blood post-ACLR.** Neutrophil and monocyte counts in whole blood after vehicle or PolySA treatment at 12hrs or 12hrs/48hrs post-ACLR with counts taken at 28d post-ACLR.

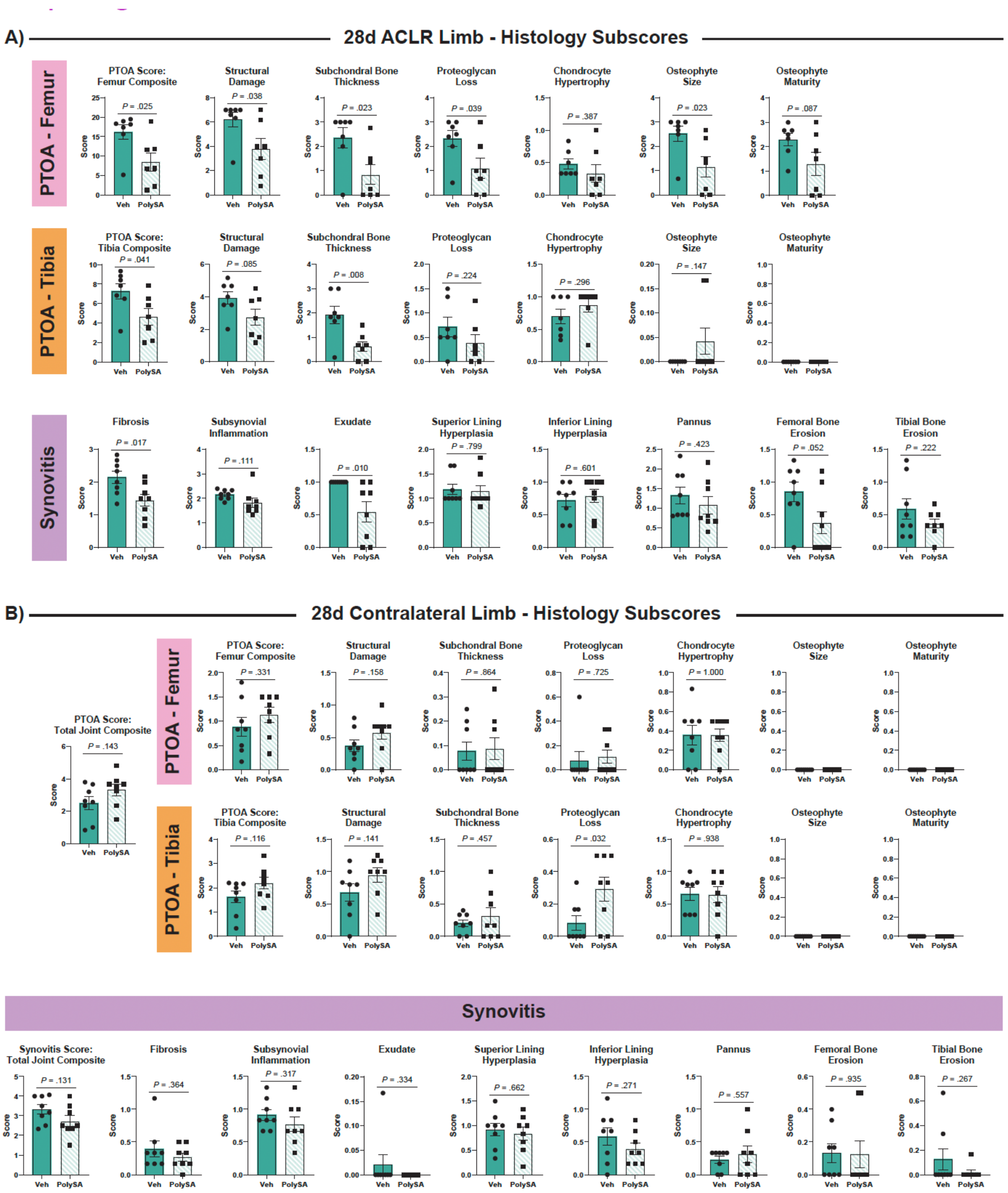


**Suppl. Fig. 11. 28d post-ACLR histopathology subscores.** Graphs show each individual subscore of PTOA and synovitis severity at 28d post-ACLR for the (A) ACLR limb and the (B) contralateral limb (n=7-8 per treatment). Graphs show the mean +/- SEM.

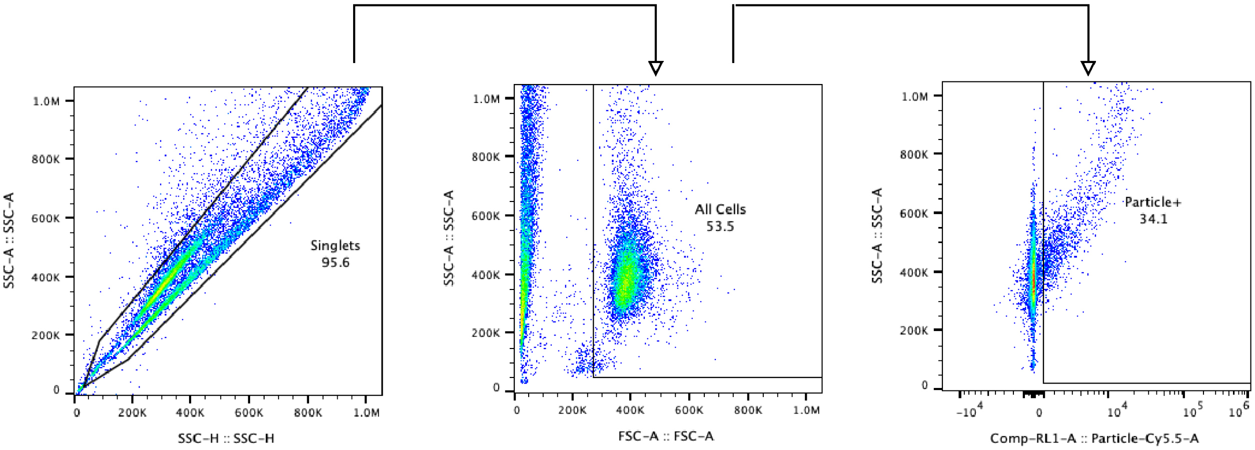


**Suppl. Fig. 12. Flow cytometry gating for human monocyte activation and uptake of PolySA particles.**

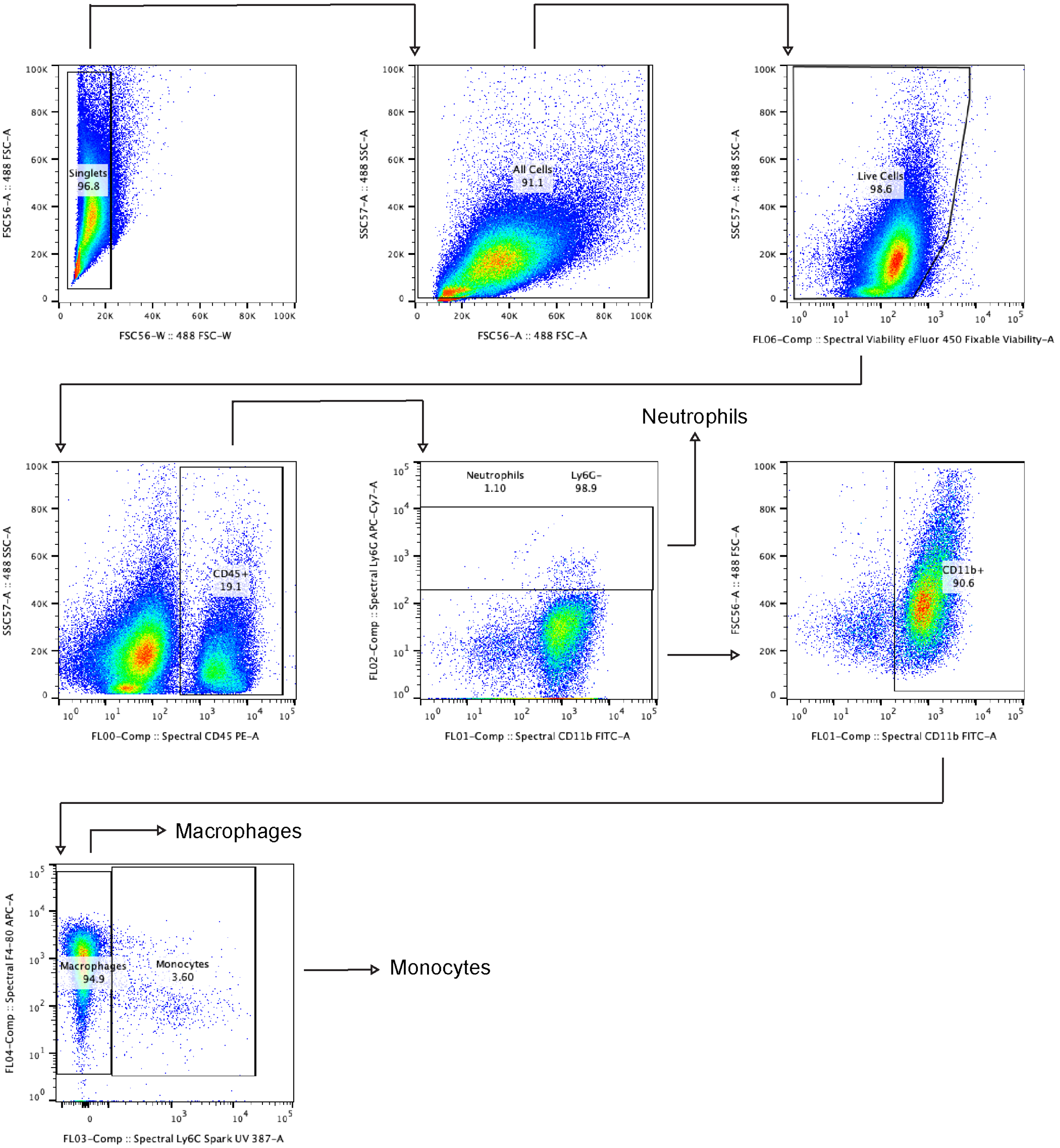


**Suppl. Fig. 13. Immune cell gating strategy for flow cytometric analysis of synovial tissue.**

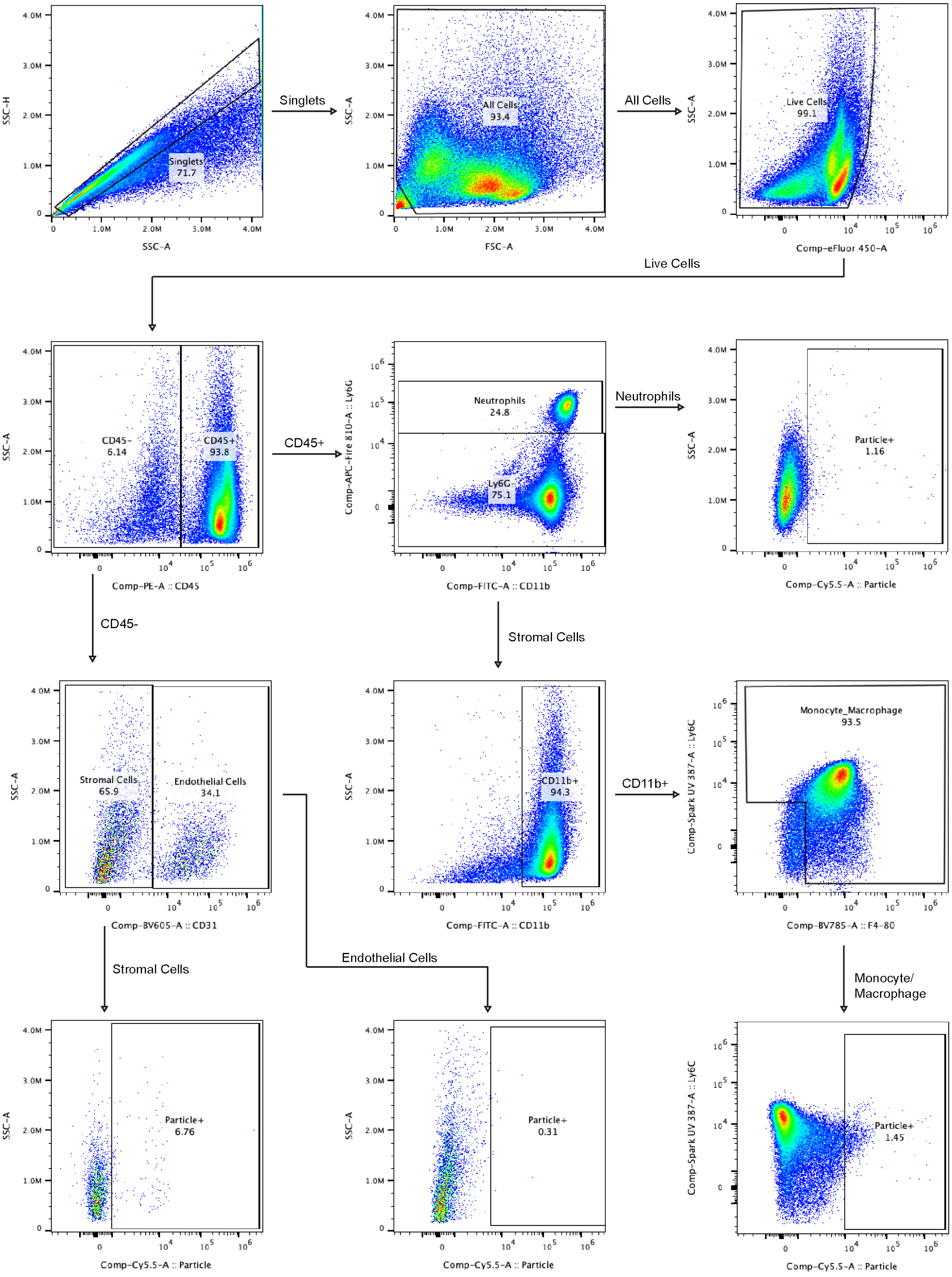


**Suppl. Fig. 14. Immune cell gating strategy for flow cytometric analysis of Cy5.5-PolySA positive particles synovial tissue.**

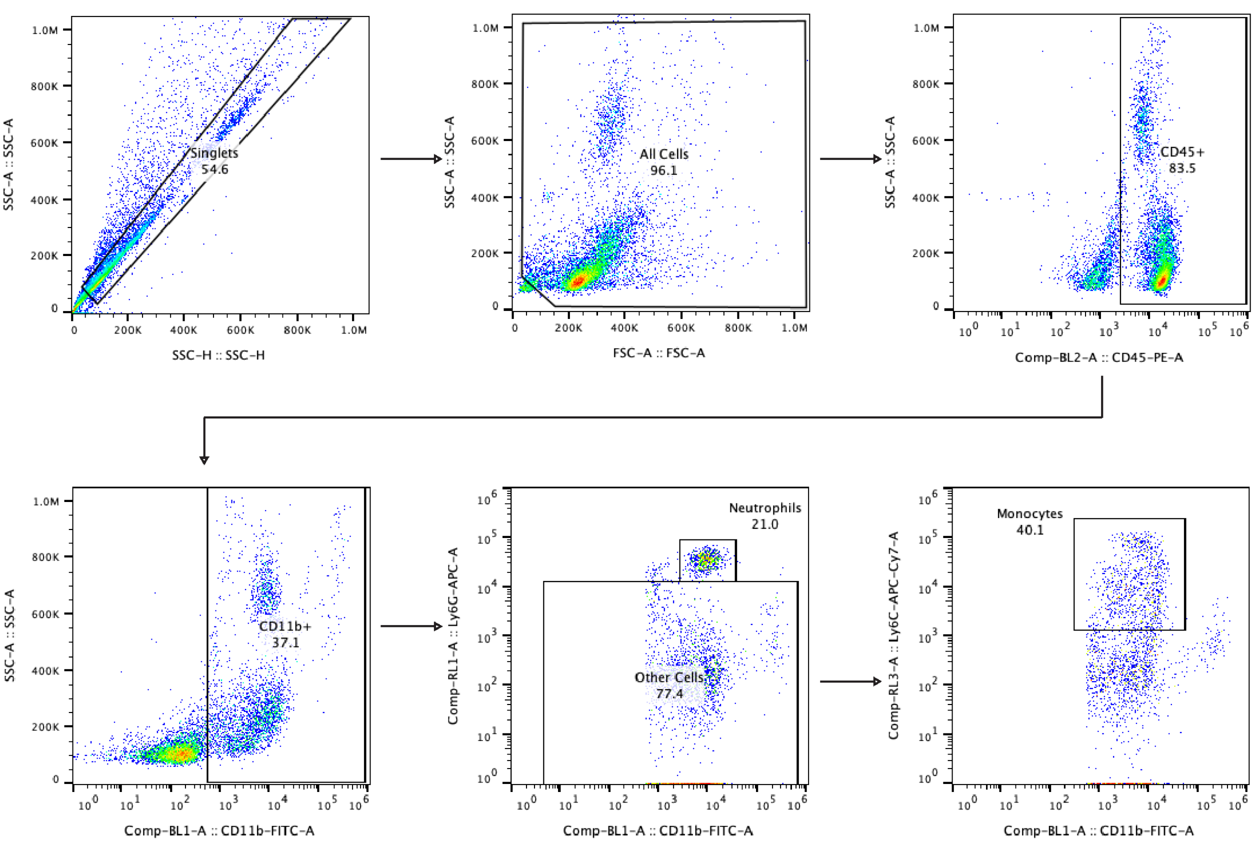


**Suppl. Fig. 15. Immune cell gating strategy for flow cytometric analysis of whole blood.**

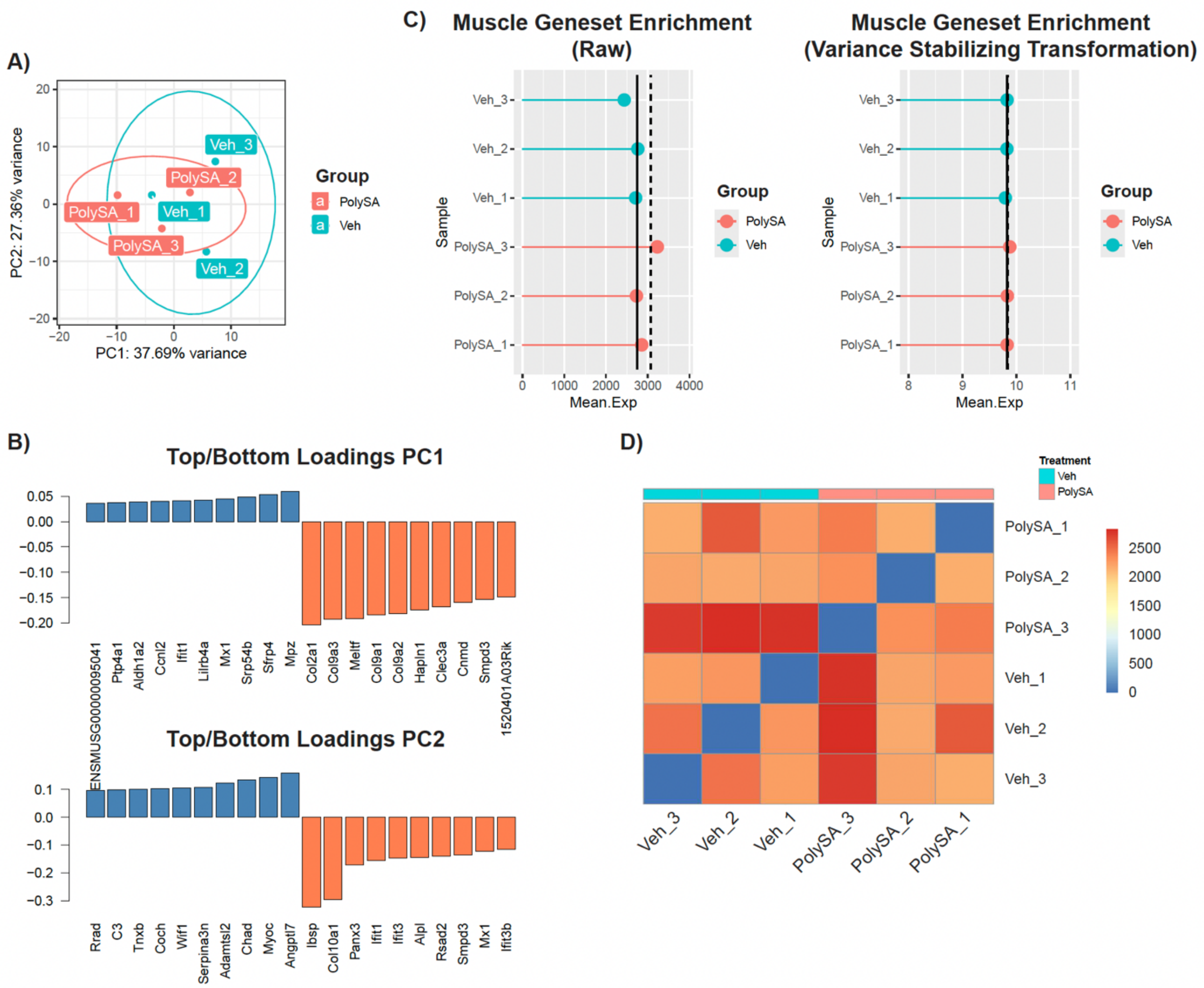


**Suppl. Fig. 16. Bulk RNAseq quality control 7d post-ACLR.** (A) Principal component analysis (PCA) plot showing 95% confidence intervals. B) Top and bottom gene loadings for principal components 1 (PC1) and 2 (PC2) from PCA analysis of normalized gene expression data. Positive values indicate genes whose expression patterns increase along the respective PC axis, while negative values indicate the converse. (C) Raw muscle gene set enrichment score and muscle gene set enrichment score with a variance stabilizing transformation. (D) Manhattan correlation plot of each individual sample.

### Supplemental tables

**Suppl. Table 1.** Positive log2FC = up in PolySA rel. to Veh. Negative log2FC = down in PolySA rel. to Veh.

| Gene | baseMean | log2FoldChange | lfcSE | stat | pvalue | padj |
|---|---|---|---|---|---|---|
| Spink5 | 94.7982043 | 1.716368991 | 0.46169794 | 3.717514951 | 0.00020119 | 0.02765405 |
| Fzd9 | 703.930929 | 1.613882645 | 0.30731085 | 5.251629293 | 1.5076E-07 | 0.00010065 |
| Hp | 1384.52879 | 1.498247342 | 0.26770432 | 5.596649749 | 2.1853E-08 | 2.5532E-05 |
| Gldn | 369.085972 | 1.367717852 | 0.22083717 | 6.193331784 | 5.8906E-10 | 1.6517E-06 |
| Slc24a2 | 77.1466061 | 1.329137009 | 0.37271515 | 3.566093308 | 0.00036234 | 0.03883941 |
| Has2 | 1165.99139 | 1.32012565 | 0.24422569 | 5.405351284 | 6.4681E-08 | 5.3343E-05 |
| Inhba | 6152.55307 | 1.270583212 | 0.21744753 | 5.843171619 | 5.1216E-09 | 9.3587E-06 |
| Dusp8 | 1424.17177 | 1.240021601 | 0.23432377 | 5.291915463 | 1.2104E-07 | 8.9316E-05 |
| Serpine1 | 9079.75349 | 1.20463549 | 0.23694199 | 5.084094675 | 3.6938E-07 | 0.00020715 |
| Gldc | 241.967372 | 1.184384474 | 0.2601951 | 4.551909249 | 5.3161E-06 | 0.00158579 |
| Sox9 | 3316.04124 | 1.168698974 | 0.23629327 | 4.945968079 | 7.5766E-07 | 0.00032189 |
| Pappa2 | 2209.80134 | 1.138617226 | 0.20293447 | 5.610762974 | 2.0144E-08 | 2.5532E-05 |
| Lrrn3 | 58.8067607 | 1.128512723 | 0.32283621 | 3.495620068 | 0.00047296 | 0.04604807 |
| Map7d2 | 81.2801025 | 1.064463144 | 0.2804337 | 3.795774689 | 0.00014718 | 0.02149487 |
| Tnfaip6 | 4656.46707 | 1.060500533 | 0.2074292 | 5.112590417 | 3.1777E-07 | 0.00018729 |
| Sox6 | 759.878895 | 0.9796691 | 0.2755433 | 3.555408912 | 0.00037739 | 0.03978219 |
| Mmp9 | 1009.00795 | 0.922036291 | 0.18536268 | 4.974228448 | 6.5508E-07 | 0.00031204 |
| Fgf7 | 965.490725 | 0.887786727 | 0.21765558 | 4.078860327 | 4.5257E-05 | 0.00893667 |
| Xylt1 | 6011.83518 | 0.869534118 | 0.22125046 | 3.930089584 | 8.4914E-05 | 0.01417259 |
| 4930523C07Rik | 1434.9675 | 0.868619098 | 0.23684308 | 3.667487788 | 0.00024495 | 0.03121938 |
| Tnn | 44716.5891 | 0.865384691 | 0.23119075 | 3.743163099 | 0.00018172 | 0.02573423 |
| Ptgs2 | 4623.56238 | 0.83449823 | 0.16644027 | 5.013799993 | 5.3365E-07 | 0.00027228 |
| Fgfr2 | 2345.28961 | 0.817739207 | 0.11478404 | 7.124154042 | 1.0472E-12 | 4.894E-09 |
| B4galnt3 | 349.348332 | 0.815255623 | 0.19707705 | 4.136735444 | 3.5228E-05 | 0.0074902 |
| Per1 | 2309.09015 | 0.809583257 | 0.22051968 | 3.671251687 | 0.00024137 | 0.03104536 |
| Nrxn2 | 319.546199 | 0.807711864 | 0.17611894 | 4.586172649 | 4.5145E-06 | 0.0014065 |
| Rgs4 | 3065.88083 | 0.761419946 | 0.10513206 | 7.242509537 | 4.4046E-13 | 3.0876E-09 |
| Cdkl5 | 286.930561 | 0.760304052 | 0.21322807 | 3.565684683 | 0.00036291 | 0.03883941 |
| Sox5 | 999.643888 | 0.738193397 | 0.15625575 | 4.724263832 | 2.3095E-06 | 0.00083024 |
| Smoc1 | 10544.0043 | 0.736432096 | 0.12329932 | 5.972718223 | 2.3333E-09 | 5.4522E-06 |
| Errfi1 | 7803.06912 | 0.734678468 | 0.18321588 | 4.009906068 | 6.0743E-05 | 0.01086877 |
| Pvr | 1528.96872 | 0.705099897 | 0.17019691 | 4.142847713 | 3.4302E-05 | 0.0074902 |
| Pde4d | 939.747005 | 0.703313049 | 0.20036257 | 3.510201832 | 0.00044777 | 0.04463366 |
| Ptx3 | 2507.7112 | 0.664060823 | 0.16037391 | 4.140703595 | 3.4624E-05 | 0.0074902 |
| Trps1 | 5734.51431 | 0.658286877 | 0.08931707 | 7.37022444 | 1.7034E-13 | 2.3882E-09 |

| | | | | | | |
|---|---|---|---|---|---|---|
| Sele | 1405.43275 | 0.653036276 | 0.1713019 | 3.812195284 | 0.00013774 | 0.02054347 |
| Fgfrl1 | 3910.19622 | 0.637285611 | 0.18136236 | 3.513880224 | 0.00044161 | 0.04461804 |
| Crlf1 | 3658.79076 | 0.60046784 | 0.09028183 | 6.651037297 | 2.9103E-11 | 1.0201E-07 |
| Trib1 | 3682.57279 | 0.586257137 | 0.16375251 | 3.580141315 | 0.00034341 | 0.03810051 |
| Rcan1 | 6055.73463 | 0.571960852 | 0.13827088 | 4.136524391 | 3.5261E-05 | 0.0074902 |
| Zbtb10 | 955.093056 | 0.504624244 | 0.12590526 | 4.007967996 | 6.1243E-05 | 0.01086877 |
| Dlc1 | 5831.04546 | 0.502750362 | 0.11223979 | 4.479252535 | 7.4905E-06 | 0.0021432 |
| Fosl2 | 11211.5544 | 0.477001309 | 0.11254824 | 4.238194183 | 2.2532E-05 | 0.00564117 |
| Filip1l | 2884.8757 | 0.398570019 | 0.10855994 | 3.671427981 | 0.0002412 | 0.03104536 |
| Lmo4 | 2035.78656 | 0.388905232 | 0.10980313 | 3.541841156 | 0.00039734 | 0.04126498 |
| Hif1a | 16018.2532 | 0.364845273 | 0.10055899 | 3.628171652 | 0.00028544 | 0.0339136 |
| Lgmn | 10861.4045 | -0.323423713 | 0.09205356 | -3.513429687 | 0.00044236 | 0.04461804 |
| Angptl1 | 11832.582 | -0.323432971 | 0.08872036 | -3.645532675 | 0.00026684 | 0.03281645 |
| Ddah2 | 2271.03061 | -0.328770621 | 0.09186534 | -3.57883216 | 0.00034513 | 0.03810051 |
| Atp13a2 | 1949.36357 | -0.374709727 | 0.10073549 | -3.719738805 | 0.00019943 | 0.02765405 |
| Ptprc | 1938.99271 | -0.37966038 | 0.09740313 | -3.897825356 | 9.706E-05 | 0.01528973 |
| Sipa1 | 1869.12565 | -0.394356643 | 0.11158815 | -3.534036776 | 0.00040926 | 0.04219032 |
| Itgb2 | 2846.91117 | -0.403597141 | 0.09451913 | -4.270004992 | 1.9547E-05 | 0.00498267 |
| Pla2g15 | 2311.84911 | -0.413339495 | 0.09336927 | -4.426932744 | 9.5583E-06 | 0.00262758 |
| Cd68 | 2934.20283 | -0.447137498 | 0.1280363 | -3.492271319 | 0.00047893 | 0.0461872 |
| Arhgap30 | 836.44703 | -0.450571068 | 0.11310701 | -3.983582042 | 6.7884E-05 | 0.01189671 |
| Cyba | 1505.89106 | -0.455874934 | 0.13099325 | -3.480140703 | 0.00050115 | 0.04778414 |
| Abca1 | 5515.14575 | -0.456107584 | 0.10852721 | -4.202702552 | 2.6375E-05 | 0.00648725 |
| Nckap1l | 2304.1243 | -0.459088449 | 0.11035633 | -4.160055353 | 3.1817E-05 | 0.0074902 |
| Pamr1 | 11401.3831 | -0.459584975 | 0.12117344 | -3.792786551 | 0.00014897 | 0.02153099 |
| Unc93b1 | 3769.67414 | -0.460651071 | 0.090131 | -5.110906278 | 3.2062E-07 | 0.00018729 |
| Ctsc | 4297.59129 | -0.464465252 | 0.11878824 | -3.910027346 | 9.2286E-05 | 0.01470279 |
| Abca9 | 3076.68884 | -0.476416447 | 0.12974997 | -3.67180392 | 0.00024084 | 0.03104536 |
| Tnmd | 21863.5814 | -0.478865994 | 0.13695459 | -3.496531171 | 0.00047135 | 0.04604807 |
| Lcp2 | 551.764139 | -0.481655387 | 0.13520713 | -3.562352066 | 0.00036755 | 0.03903795 |
| Laptm5 | 6185.43252 | -0.486373036 | 0.09039187 | -5.38071665 | 7.419E-08 | 5.7786E-05 |
| Inpp5d | 1737.32052 | -0.499604071 | 0.1102013 | -4.533558977 | 5.7998E-06 | 0.00169403 |
| Tmem86a | 1337.39391 | -0.504446699 | 0.1255575 | -4.017654915 | 5.878E-05 | 0.01070258 |
| Arhgdib | 1032.72339 | -0.50549499 | 0.13837094 | -3.653187638 | 0.000259 | 0.03242185 |
| Lrrc25 | 461.282297 | -0.506686698 | 0.14150154 | -3.58078582 | 0.00034256 | 0.03810051 |
| Rps6ka1 | 724.07861 | -0.513940801 | 0.13391453 | -3.837827098 | 0.00012413 | 0.0187126 |
| Ncf1 | 989.259906 | -0.514171529 | 0.13022132 | -3.948443484 | 7.8661E-05 | 0.01344911 |
| Ctss | 5736.72212 | -0.51561601 | 0.12456924 | -4.139192155 | 3.4853E-05 | 0.0074902 |
| Lpcat2 | 486.35837 | -0.52143319 | 0.14857596 | -3.509539479 | 0.00044888 | 0.04463366 |
| Irf5 | 733.107996 | -0.523451697 | 0.14826994 | -3.530396569 | 0.00041494 | 0.04246292 |
| Csf1r | 8829.12912 | -0.526181382 | 0.13343515 | -3.943349223 | 8.0352E-05 | 0.01357263 |

| | | | | | | |
|---|---|---|---|---|---|---|
| Ccr5 | 934.360717 | -0.532292889 | 0.13989083 | -3.805059266 | 0.00014177 | 0.0209223 |
| Cfp | 854.292441 | -0.534228535 | 0.11212491 | -4.764583651 | 1.8924E-06 | 0.00071708 |
| Fcgr2b | 1860.00112 | -0.534232454 | 0.11263135 | -4.743194966 | 2.1037E-06 | 0.00077617 |
| Rassf2 | 2784.20118 | -0.534452552 | 0.14704327 | -3.634661713 | 0.00027835 | 0.03364143 |
| Arhgap45 | 1247.61637 | -0.544678906 | 0.12694913 | -4.290528755 | 1.7825E-05 | 0.00471517 |
| Syk | 1275.17525 | -0.545568522 | 0.11232274 | -4.857151135 | 1.1909E-06 | 0.00049106 |
| Stab1 | 11124.6627 | -0.558983189 | 0.15120775 | -3.696789295 | 0.00021834 | 0.02915404 |
| Trf | 2136.92549 | -0.562503751 | 0.11107337 | -5.064254097 | 4.1E-07 | 0.00022109 |
| Cd300ld | 523.980711 | -0.567134006 | 0.14096946 | -4.02309828 | 5.7438E-05 | 0.01059571 |
| Anxa3 | 1366.57034 | -0.576916534 | 0.15951848 | -3.616612544 | 0.00029848 | 0.03516588 |
| Fcer1g | 1252.42459 | -0.584385619 | 0.13678293 | -4.272357864 | 1.9342E-05 | 0.00498267 |
| C1qc | 4819.12887 | -0.586389726 | 0.14108382 | -4.156321646 | 3.2341E-05 | 0.0074902 |
| Pirb | 900.175049 | -0.593688968 | 0.13472823 | -4.406566962 | 1.0502E-05 | 0.00283155 |
| Mbp | 609.923963 | -0.595683022 | 0.15428285 | -3.860980059 | 0.00011293 | 0.01721003 |
| Adgre1 | 4474.104 | -0.605133024 | 0.11508133 | -5.258307603 | 1.4539E-07 | 0.00010065 |
| Abcc3 | 640.994189 | -0.607494274 | 0.17464787 | -3.478395022 | 0.00050443 | 0.04778414 |
| Otulinl | 554.686259 | -0.615336044 | 0.16940009 | -3.632442242 | 0.00028075 | 0.03364218 |
| Pld4 | 1586.7149 | -0.624565906 | 0.11248763 | -5.552307391 | 2.8192E-08 | 3.0404E-05 |
| Siglec1 | 1268.75385 | -0.628491185 | 0.13719985 | -4.580844715 | 4.631E-06 | 0.00141145 |
| Cd300c2 | 357.77488 | -0.63138329 | 0.16885054 | -3.739302851 | 0.00018453 | 0.02587129 |
| Mpeg1 | 6290.11309 | -0.635539645 | 0.12818465 | -4.958001067 | 7.1222E-07 | 0.00031204 |
| C3ar1 | 2119.2853 | -0.643665453 | 0.1739112 | -3.701115683 | 0.00021465 | 0.02893695 |
| Snx20 | 232.722184 | -0.648122609 | 0.18011678 | -3.598346578 | 0.00032025 | 0.03710627 |
| Cd53 | 1064.35384 | -0.648665479 | 0.13995821 | -4.634708336 | 3.5744E-06 | 0.00116542 |
| Fcrls | 2776.84041 | -0.655183036 | 0.16724501 | -3.917504191 | 8.947E-05 | 0.01458577 |
| Itgax | 455.591102 | -0.657100992 | 0.18350933 | -3.580749817 | 0.00034261 | 0.03810051 |
| C1qa | 6068.16603 | -0.65935793 | 0.13788472 | -4.781950571 | 1.736E-06 | 0.00067608 |
| Cybb | 2595.0273 | -0.667669547 | 0.11440123 | -5.83620931 | 5.3402E-09 | 9.3587E-06 |
| Ms4a7 | 1139.4514 | -0.669869952 | 0.16558247 | -4.045536546 | 5.2203E-05 | 0.00984817 |
| Themis2 | 500.789518 | -0.670824392 | 0.14543185 | -4.612637369 | 3.9759E-06 | 0.00126687 |
| Cd300a | 444.033181 | -0.682225478 | 0.19215067 | -3.550471509 | 0.00038454 | 0.0402334 |
| Slamf7 | 237.781388 | -0.68377645 | 0.17652366 | -3.87356821 | 0.00010725 | 0.01652409 |
| Clec4a1 | 732.764239 | -0.687332204 | 0.16572205 | -4.147500069 | 3.3613E-05 | 0.0074902 |
| Chst5 | 388.603808 | -0.693315014 | 0.17717935 | -3.913069058 | 9.113E-05 | 0.01468562 |
| Bbc3 | 223.123608 | -0.695678339 | 0.18431854 | -3.774326411 | 0.00016044 | 0.02295283 |
| C1qb | 6201.21186 | -0.699450638 | 0.14086344 | -4.96545196 | 6.8541E-07 | 0.00031204 |
| Tlr13 | 677.799472 | -0.700957727 | 0.14901332 | -4.7039938 | 2.5512E-06 | 0.00087472 |
| Il10ra | 942.429795 | -0.713715704 | 0.17558494 | -4.064788902 | 4.8076E-05 | 0.00936146 |
| Card6 | 182.68029 | -0.717077514 | 0.20539974 | -3.491131544 | 0.00048098 | 0.0461872 |
| Trem2 | 1038.18074 | -0.717303684 | 0.17571741 | -4.082143433 | 4.4622E-05 | 0.00893667 |
| Ccl4 | 373.676251 | -0.724102761 | 0.20218143 | -3.581450301 | 0.00034169 | 0.03810051 |

| | | | | | | |
|---|---|---|---|---|---|---|
| Txnip | 7024.52088 | -0.745380198 | 0.12985577 | -5.740062335 | 9.4642E-09 | 1.4743E-05 |
| Clec4a3 | 453.805699 | -0.751384774 | 0.18356387 | -4.093315132 | 4.2525E-05 | 0.00864057 |
| Irf8 | 705.703643 | -0.755637641 | 0.18416699 | -4.103002643 | 4.0782E-05 | 0.00840834 |
| Fcgr1 | 468.252161 | -0.760910353 | 0.15347073 | -4.958016258 | 7.1217E-07 | 0.00031204 |
| Selenop | 10530.5723 | -0.766698962 | 0.17215439 | -4.4535546 | 8.446E-06 | 0.00236826 |
| Ifi27l2a | 234.30698 | -0.768341397 | 0.21420772 | -3.586898696 | 0.00033463 | 0.03810051 |
| Gatm | 250.826491 | -0.778009041 | 0.21103242 | -3.686680112 | 0.0002272 | 0.03005024 |
| Apoe | 32345.0863 | -0.779987672 | 0.14248538 | -5.474159507 | 4.3959E-08 | 3.8519E-05 |
| Lyz2 | 27077.2851 | -0.780035592 | 0.1388151 | -5.61924175 | 1.918E-08 | 2.5532E-05 |
| Ms4a6c | 396.018229 | -0.784246813 | 0.14959168 | -5.242583218 | 1.5834E-07 | 0.00010091 |
| Adamts15 | 2478.86964 | -0.798469814 | 0.1974751 | -4.04339485 | 5.2683E-05 | 0.00984817 |
| Tifab | 306.503422 | -0.800212153 | 0.19331346 | -4.139453977 | 3.4813E-05 | 0.0074902 |
| Ptafr | 458.586984 | -0.803905711 | 0.19513075 | -4.119830958 | 3.7915E-05 | 0.00793386 |
| Sash3 | 316.681843 | -0.806790672 | 0.14644488 | -5.509176331 | 3.6052E-08 | 3.4908E-05 |
| Npl | 164.905209 | -0.841954218 | 0.23041132 | -3.654135607 | 0.00025805 | 0.03242185 |
| Tnfaip8l2 | 215.312567 | -0.844501328 | 0.1745567 | -4.837977106 | 1.3117E-06 | 0.00052542 |
| Gadd45a | 655.71947 | -0.854285177 | 0.23697275 | -3.604993345 | 0.00031216 | 0.0364708 |
| Gpr65 | 161.54251 | -0.858565625 | 0.24018974 | -3.574530876 | 0.00035086 | 0.03842974 |
| Clec7a | 345.396697 | -0.866746583 | 0.21428557 | -4.044820031 | 5.2363E-05 | 0.00984817 |
| Ly86 | 544.47237 | -0.873030252 | 0.18795365 | -4.644923132 | 3.402E-06 | 0.00113563 |
| Cx3cr1 | 2010.7325 | -0.876511517 | 0.24103884 | -3.63639122 | 0.00027648 | 0.03364143 |
| Cd4 | 212.815755 | -0.891394321 | 0.22714698 | -3.924306357 | 8.698E-05 | 0.01434658 |
| Dact2 | 251.318836 | -0.9007492 | 0.17978378 | -5.010180684 | 5.4379E-07 | 0.00027228 |
| P2ry13 | 130.420525 | -0.903394219 | 0.23231665 | -3.888633106 | 0.00010081 | 0.01570402 |
| N4bp2l1 | 135.61073 | -0.937788795 | 0.25318484 | -3.703969021 | 0.00021225 | 0.02889103 |
| Hic1 | 1404.24044 | -0.965077855 | 0.27033159 | -3.569978076 | 0.00035701 | 0.03880074 |
| Cd274 | 134.930949 | -1.019139387 | 0.27922106 | -3.649937435 | 0.0002623 | 0.03254429 |
| Dpp4 | 536.142335 | -1.079261359 | 0.27193419 | -3.968832934 | 7.2225E-05 | 0.01250125 |
| Fgf16 | 93.1582143 | -1.282492088 | 0.36599623 | -3.504112805 | 0.00045813 | 0.0452324 |
| Ebi3 | 72.6451597 | -1.440359538 | 0.30623471 | -4.703449655 | 2.558E-06 | 0.00087472 |
| Wnt2 | 109.2646 | -1.858565765 | 0.33773958 | -5.502955086 | 3.7348E-08 | 3.4908E-05 |

### Suppl. Table 2. Human flow cytometry antibodies.

| Marker | Fluorophore | Manufacturer | Catalog # |
|---|---|---|---|
| Human FcX | NA | BioLegend | 422302 |
| Particle | Cy5.5 | NA | NA |
| CD14 | PerCP | BioLegend | 325632 |
| CD62L | BV421 | BioLegend | 304828 |
| CD11b | AF488 | BioLegend | 301318 |

### Suppl. Table 3. Mouse flow cytometry antibodies.

| Synovial Tissue | | | |
|---|---|---|---|
| **Marker** | **Fluorophore** | **Manufacturer** | **Catalog #** |
| Mouse FcX | NA | BioLegend | 101320 |
| Viability | eFluor 450 | eBioscience | 65086318 |
| CD45 | PE | BioLegend | 103106 |
| CD11b | FITC | BioLegend | 101206 |
| Ly6G | APC/Cy7 | BioLegend | 127624 |
| Ly6C | Spark UV 387 | BioLegend | 128060 |
| F4/80 | APC | BioLegend | 123116 |
| CD31 | SB 610 | Bio-Rad | MCA2388SB610 |

| Whole Blood | | | |
|---|---|---|---|
| **Marker** | **Fluorophore** | **Manufacturer** | **Catalog #** |
| Mouse FcX | NA | BioLegend | 101320 |
| CD45 | PE | BioLegend | 103106 |
| CD11b | FITC | BioLegend | 101206 |
| Ly6G | APC | BioLegend | 127614 |
| Ly6C | APC-Cy7 | BioLegend | 128026 |

### Suppl. Table 4. Histopathologic PTOA severity scoring.

| **Category**<br>**Range:**<br>**Region(s):** | **0** | **1** | **2** | **3** | **4** | **5** | **6** | **7** |
|---|---|---|---|---|---|---|---|---|
| **Structural damage**<br>**Range:** 0-7<br>**Region(s):**<br>1. Femur<br>2. Tibia | Normal cartilage. | Roughened surface with small fibrillations, wavy articular surface. | Fibrillations immediately below superficial layer or some loss of laminal surface. | Horizontal cracks or separations between calcified and non-calcified cartilage. | Mild loss of non-calcified cartilage (<10% surface area). | Moderate loss of non-calcified cartilage (10-50% surface area). | Severe loss of non-calcified cartilage (>50% surface area). | Erosion of cartilage to subchondral bone (any percent surface area). |
| **Proteoglycan loss**<br>**Range:** 0-3<br>**Region(s):**<br>1. Femur<br>2. Tibia | Normal cartilage. | Decreased but not complete loss of Safranin-O staining in non-calcified areas. Saf-O loss does not penetrate completely from superficial to deep cartilage. | Saf-O loss penetrates completely from superficial to deep cartilage. Focal loss of Safranin-O staining in non-calcified area (<30% surface area). | Saf-O loss penetrates completely from superficial to deep cartilage. Diffuse loss of Safranin-O staining in non-calcified cartilage (>30% surface area). | | | | |
| **Chondrocyte hypertrophy**<br>**Range:** 0-1<br>**Region(s):**<br>1. Femur<br>2. Tibia | None. | Enlarged chondrocyte lacunae with lack of Saf-O stain around collapsed cell. | | | | | | |
| **Osteophyte size**<br>**Range:** 0-3<br>**Region(s):**<br>1. Femur<br>2. Tibia | None. | Small – ≤1x thickness as adjacent cartilage. | Medium – >1x to 3x as thick as adjacent cartilage. | Large - >3x thicker than adjacent cartilage. | | | | |
| **Osteophyte maturity**<br>**Range:** 0-3<br>**Region(s):**<br>1. Femur<br>2. Tibia | None. | Predominately cartilage. | Mixed cartilage and bone with vascular invasion. | Predominately bone. | | | | |
| **Subchondral bone thickening**<br>**Range:** 0-3<br>**Region(s):**<br>1. Femur<br>2. Tibia | Normal SCB. | Mild thickening, <50% increase. | Moderate thickening, 50-100% increase. | Severe thickening, >100% increase. | | | | |

### Suppl. Table 5. Histopathologic synovitis severity scoring.

| **Category**<br>**Range:**<br>**Region(s):** | **0** | **1** | **2** | **3** |
|---|---|---|---|---|
| **Pannus**<br>**Range:** 0-3<br>**Region(s):**<br>1. Anterior synovium/tibia | None. | Mild: Pannus has migrated onto bone but is not encroaching on articular surface. It is in the transition zone where there is calcified cartilage, but you are not yet at the articular surface. | Moderate: Pannus has migrated through the transition zone and <1x cartilage depth onto the articular surface. | Severe: Pannus has migrated > 1x cartilage depth onto the articular surface. |
| **Bone erosion**<br>**Range:** 0-3<br>**Region(s):**<br>1. Anterior femur<br>2. Anterior tibia | None. | Partial thickness loss of cortical bone only. Wavy surface. | Focal complete loss of cortical bone - communication with marrow cavity at one small vascular communication site. A single large “offshoot” of cortical bone erosion. | Widespread complete loss of cortical bone - communication with marrow cavity at multiple sites or broad area loss of cortical bone. |
| **Synovial lining hyperplasia**<br>**Range:** 0-3<br>**Region(s):**<br>1. Anterior, superior synovium<br>2. Anterior, inferior synovium | 1 cell thick. | Mild: 2-3 cells thick. | Moderate: 4-5 cells thick. | Severe: ≥6 cells thick. |
| **Subsynovial inflammation**<br>**Range:** 0-3<br>**Region(s):**<br>1. Anterior synovium | None. | One pocket of densely associated inflammatory cells. | Two to three pockets of densely associated inflammatory cells. These are focal areas of dense subsynovial WBC infiltrate – but still predominantly normal subsynovial areolar connective tissue present. | ≥ Four pockets of densely associated inflammatory cells. This is widespread dense subsynovial WBC infiltrate with markedly reduced or little/no normal areolar connective tissue evident and some lymphoid follicle formation. |
| **Synovial fibrosis**<br>**Range:** 0-3<br>**Region(s):**<br>1. Anterior synovium | None (less than 10% to account for the immediate sublining and regular matrix around blood vessels). | Dispersed fibrosis (10% to 1/3 of synovial area). | Moderate fibrosis (>1/3 to 2/3 of synovial area). | Severe fibrosis (>2/3 of synovial area). |
| **Synovial exudate**<br>**Range:** 0-1<br>**Region(s):**<br>1. Anterior synovium | None. | Infiltration of inflammatory cells (neutrophils, macrophages, and/or lymphocytes) or fibrin in the synovial cavity. | | |
